# Actin-related protein Alp1 governs malaria parasite motility and transmission

**DOI:** 10.64898/2026.07.14.737759

**Authors:** Yukino Kobayashi, Caroline Busse, Annika Binder, Amélie Moll, Friedrich Frischknecht, Ross G. Douglas

## Abstract

Motility of the malaria-causing parasite *Plasmodium* is essential for transmission to and from mosquitoes, with the turnover of actin filaments being a central feature of productive cell movement. Actin-related proteins (Arps) are known to play critical roles in motility, trafficking and chromatin remodelling. Here, we show that the actin-like protein 1 (Alp1), an apicomplexan Arp, is essential for *Plasmodium* ookinete motility and establishing of infection in mosquitoes. We identified an insertion region in subdomain 4 that contributes to Alp1 function in ookinetes and show novel actin filament structures in ookinetes. A combination of gene deletion and actin filament recognizing chromobody expression revealed a role of Alp1 in promoting actin filament turnover in ookinetes. We have thus identified a novel Arp that has evolved a specialist function to regulate actin dynamics, govern malaria parasite motility and facilitate malaria transmission.

## Introduction

The malaria causing parasite *Plasmodium* must adapt, replicate and spread in a variety of tissue types of two different hosts (vertebrate and mosquito)^1^. Upon ingestion of infected blood by an *Anopheles* mosquito, male parasites rapidly develop into motile gametes that fertilize a female^2^. The zygotes then develop into motile ookinetes that actively penetrate the midgut epithelium to form an oocyst^3^. Within oocysts sporozoites develop that penetrate the salivary gland and are transmitted during a bite using active motility to migrate in the skin and enter hepatocytes^4^. Cell motility is therefore essential for parasite transmission to and from mosquitoes^5^. Ookinetes and sporozoites move via an uncommon mode of locomotion termed gliding motility, whereby the parasite moves without any obvious changes in cell shape^6,7^. Ookinetes move at high speeds (3-5 microns/min) comparable to neutrophils, while sporozoites are 10-fold faster. Actin is a central molecule in gliding, whereby a myosin motor powerstroke results in the rearward translocation of actin filaments and actin associated membrane spanning adhesins, ultimately propelling the organism forward^8,9^. *Plasmodium* actin, like opisthokont actins, has a conserved actin core structure with four subdomains^10^ yet is highly sequence divergent from these classical actins^11^. Exchange of highly divergent subdomain regions to classical counterparts revealed the importance of subdomains 2 and 3 in parasite viability and mosquito colonization^12,13^. Exchange of the entire subdomain 4 had a pronounced effect on smooth motility and salivary gland invasion of sporozoites^12^. Some of the other key components of the gliding machinery have been identified based on homology, such as actin-binding proteins^14^. However, many divergent and highly specialized molecules that are critical for gliding motility and mosquito transmission remain unknown.

Actin-related proteins are highly sequence divergent proteins in comparison to actin and fulfill multiple essential roles in opisthokont cells^15,16^. For example Arp1 can oligomerise and, along with Arp10 and Arp11, is a critical factor in facilitating intracellular trafficking^17–19^. Arp 2 and Arp3 are famously involved in the regulation of actin and mediating branched filament networks^20–23^, while Arps 4-9 are classically referred to as nuclear Arps and facilitate chromatin dynamics^24–26^. Arps share structural similarity to actin by all having an actin fold. However, Arps have striking insertions and deletions beyond the actin fold core^15,16^. Interestingly, *Plasmodium* has decreased numbers of classically assigned Arps, with only Arp1, Arp 4 and Arp6 phylogenetically clustering with their respective classical homologues^27^. Instead, the *Plasmodium* genome contains “actin-like proteins” (Alps): Arps that do not share strong sequence homology to classical opisthokont Arps and are unique to apicomplexan parasites^27^. Very little is known about the roles of these Alps in apicomplexan parasites and only recently two Alps were reassigned as *Plasmodium* Arp2 and Arp3^28^. In *Toxoplasma*, the Alp1 homologue was refractory to gene deletion while overexpression negatively impacted cell division^29,30^. Here, we characterized the role of Alp1 in *Plasmodium*. We identified an essential role of Alp1 in transmission to mosquitoes, specifically in governing ookinete motility, together with an insertion region that contributes to Alp1 function in ookinetes. We further made use of an actin chromobody probe and show novel actin filament structures in ookinetes that are different from those found in sporozoites, revealing the similarities and differences in actin dynamics at different stages of the parasite life cycle. Finally, we show that deletion of *alp1* renders actin filaments more prone to stabilization *in vivo*, pointing to a potential role of Alp1 in promoting actin filament turnover in ookinetes. We have thus identified a unique Arp that has evolved a specialist function to regulate actin dynamics, govern ookinete motility and ultimately facilitate malaria transmission.

## Results

### Alp1 is structurally related to actin and plays an essential role in transmission to mosquitoes via ookinete motility

Alp1 is a highly sequence divergent Arp that displays 34% amino acid sequence identity to conventional actin and only 35% to the predicted closest related canonical Arp (Arp1)^27,30^. Nonetheless, the predicted structure of Alp1 displays high structural homology to the conventional actin structure (**Figure 1A**). *Alp1* transcripts are most abundant in the trophozoite, male gametocyte and liver stages^31,32^, suggesting potentially important roles in several parts of the parasite life cycle. In order to determine the contribution of Alp1 to parasite life cycle progression, we deleted the *alp1* gene (PlasmoDB gene ID: PBANKA_0936900) in the *P. berghei* rodent model of infection (**Figure 1B**) using the PlasmoGEM knockout construct (**Figure S1**, line name: PbAlp1 KO-PG)^33,34^ and characterized the ability of the knockout line to grow asexually in the blood. PbAlp1 KO-PG parasites grew at modestly slower growth rates in comparison to the complementation control (where the knockout was complemented with wild-type *P. berghei alp1,* line name: PbAlp1 Comp.) and previously published wild-type lines (**Figure 1C**, **Figure S2**)^12,35^, consistent with the growth rates observed in the PlasmoGEM screen^33,34^. The design of the original vector left 36% of the open reading frame on the 3’ end (**Figure S1**), which might facilitate blood stage growth. In order to assess whether the remnant *alp1* significantly contributes to blood stage growth, we performed a second transfection in which all known *alp1* sequence was deleted (**Figure S1**, line name: PbAlp1 KO-full). This line closely phenocopied the PbAlp1 KO-PG line indicating that the remnant sequence has no major consequence on blood stage growth (**Figure 1C**).

**Figure 1.**
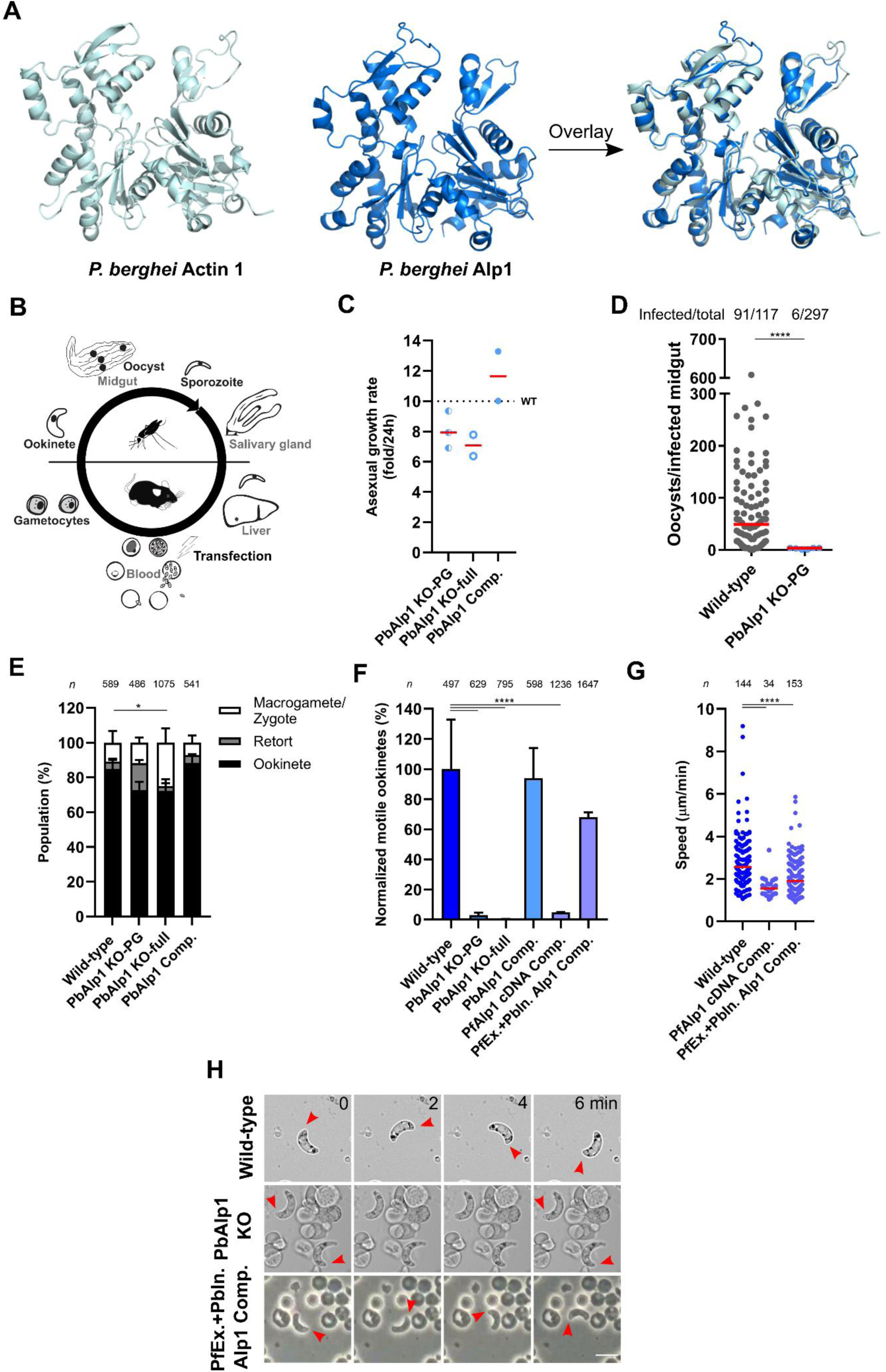
*Plasmodium* Alp1 is a highly divergent, apicomplexan unique actin-related protein essential for ookinete motility and mosquito transmission. (A) AlphaFold2 predicted structures of Alp1 (dark blue) and actin (silver). Overlay of the two structures shows the high degree of predicted structural similarity between the two proteins. (B) Schematic representation of the *Plasmodium* life cycle showing relevant stages. Transfection and selection of genetic modifications are performed with asexual blood stages. (C) Partial (KO-PG) and complete (KO-full) deletion of *alp1* resulted in only a modest effect of the blood stage growth. Dotted line indicates a published wild-type value^35^. (D) Alp1 KO parasites were fed to mosquitoes but showed a drastic reduction in both mosquito infectivity and oocyst loads. Mann-Whitney test, **** p<0.0001. (E) Quantification of relevant developmental stages revealed that ookinete maturation was not majorly affected by the absence of Alp1. Fisher’s exact test, * p<0.05. (F) Percentage representation of motile ookinete populations in each Alp1 KO and complementation line, normalized to the wild-type population. Alp1 KOs led to a drastic reduction in the motile ookinete population, indicating a critical contribution of Alp1 to ookinete motility. Cross-species complementation with *P. falciparum* (Pf) cDNA (PfAlp1 cDNA Comp.) resulted in a KO-like phenotype. In contrast, complementation of a hybrid Alp1 consisting of Pf exons and Pb introns (PfEx.+PbIn. Alp1 Comp.), significantly rescued ookinete motility, indicating that Alp1 function is conserved with the human infecting species *P. falciparum* and the importance of *P. berghei* introns. Fisher’s exact test performed on unnormalized percentage values with each corresponding wild-type control, **** p<0.0001. (G) Gliding speeds of Alp1 complementation ookinetes with or without Pb-introns show a minor decrease in speed likely due to experimental variation. Red line indicates median value. Mann-Whitney test, **** p<0.0001. (H) Representative images of wild-type, Alp1 KO-full and PfEx.+PbIn. Alp1 Comp. ookinetes acquired at 0, 2, 4 and 6 minutes during the motility assay. Wild-type and PfEx.+PbIn. Alp1 Comp. ookinetes were motile, whereas Alp1 KO-full ookinetes remained stationary. Red arrow indicates the front of the ookinete. Scale bar: 10 μm.

Malaria parasites are transmitted from one vertebrate to another by the bite of an infected mosquito. We next assessed the ability of *alp1* knockout parasites to transmit to the mosquito host. Mosquitoes were infected with either knockout or wild-type parasites and the ability of the parasite to colonize mosquitoes was determined by both the number of infected mosquitoes and the total amount of oocysts per infected mosquito midgut. Wild-type parasites could infect the large majority of mosquitoes (∼77%) with high oocyst loads (median: ∼50 oocysts per mosquito). However, *alp1* deletion resulted in a nearly complete block in transmission, whereby only 6 out of 297 mosquitoes (∼2%) were infected by knockout parasites (**Figure 1D**). These infected mosquitoes also had remarkably low oocyst loads of less than 5 per mosquito, together indicating that Alp1 is a critical transmission factor for mosquito infection.

Several developmental steps are required for the malaria parasite to form oocysts in the mosquito. After uptake in a mosquito blood meal, sexual forms (gametocytes) are activated and male gametes fertilize a female to form a zygote^2^. Over the course of 21 hours, zygotes transform first into intermediate retort forms before developing into mature ookinetes^36^. Motile ookinetes penetrate the midgut epithelium to ultimately form oocysts^3^. Knockout parasites developed into ookinetes in similar proportions to controls (**Figure 1E**), indicating no major developmental issues between blood stages and ookinetes. We next explored whether ookinetes from the knockout lines were motile. While ookinetes from control lines (wild-type and PbAlp1 Comp.) moved readily, ookinetes from knockout lines were completely immotile (**Figure 1F, H**) revealing a central role for Alp1 in ookinete motility.

### Alp1 function is conserved in a human infecting species

The rodent infecting *P. berghei* model is a powerful model to study all stages of the *Plasmodium* life cycle. To test whether Alp1 function is conserved between a rodent and human infecting species, the *alp1* knockout line was complemented with the *P. falciparum* homologue (PlasmoDB gene ID: PF3D7_1110700). We first complemented the knockout line with *P. falciparum* cDNA (**Figure S3**, line name: PfAlp1 cDNA Comp.) and assessed ookinete motility of this line. Surprisingly, complementation with *P. falciparum* cDNA produced very few motile ookinetes and thus did not rescue the knockout motility phenotype, although the few moving ookinetes could move at speeds that were only slightly slower yet in a similar range to the wild-type control (**Figure 1F-G**). The gene structure for both *P. berghei* and *P. falciparum alp1* genes are similar, containing four exons and three introns. Given that the Alp1 protein sequence is relatively highly conserved between these species (∼76% amino acid sequence identity), we hypothesized that the *P. berghei* introns might play a role in regulating expression of *alp1* needed for ookinete motility. To test this, we generated and transfected another complementation construct containing *P. falciparum alp1* exons with the corresponding *P. berghei alp1* introns (**Figure S4**, line name: PfEx.+PbIn. Alp1 Comp.). This would ensure the presence of native *P. berghei* genome elements, while producing *P. falciparum* Alp1 protein. Complementation with this construct resulted in near-complete rescue of ookinete motility (**Figure 1F-H**), demonstrating the functional conservation of Alp1 between these species.

### A subdomain 4 region on Alp1 contributes to ookinete motility

The analysis so far revealed that Alp1 is a functionally conserved transmission factor essential for ookinete motility. We next sought to characterize the structure-function relationships of Alp1 in comparison to conventional actin, the archetypal member of the actin superfamily. The predicted three-dimensional structure of Alp1 is highly similar to that of *P. berghei* actin 1 despite low sequence identity, yet comparison of primary structures indicated the presence of small insertions and deletions compared to the canonical backbone (**Figure 2A-B**, **Figure S5**). These changes were often in regions that map to the surface of the protein and were consistent between multiple *Plasmodium* species (**Figure S5**). We earmarked three regions based on size and location of the insertion or deletion: 1) there was a noticeable decrease in the size of the D-loop in subdomain 2 (residues 43-48 in comparison to actin) and in 2) an external helix of subdomain 4 (residues 229-232). Further, 3) we noticed a region of insertion on the “pointed end” of subdomain 4 (residues 251-252 in comparison to actin) that is flanked between residue sequences which are highly conserved between Alp1 and actin (“LPDG” and “PEALF”), strongly suggesting an insertion in this region (**Figure S5**). We hypothesized that, since these regions are conserved and map to the protein surface, these regions could be major contributors to specific Alp1 function.

**Figure 2.**
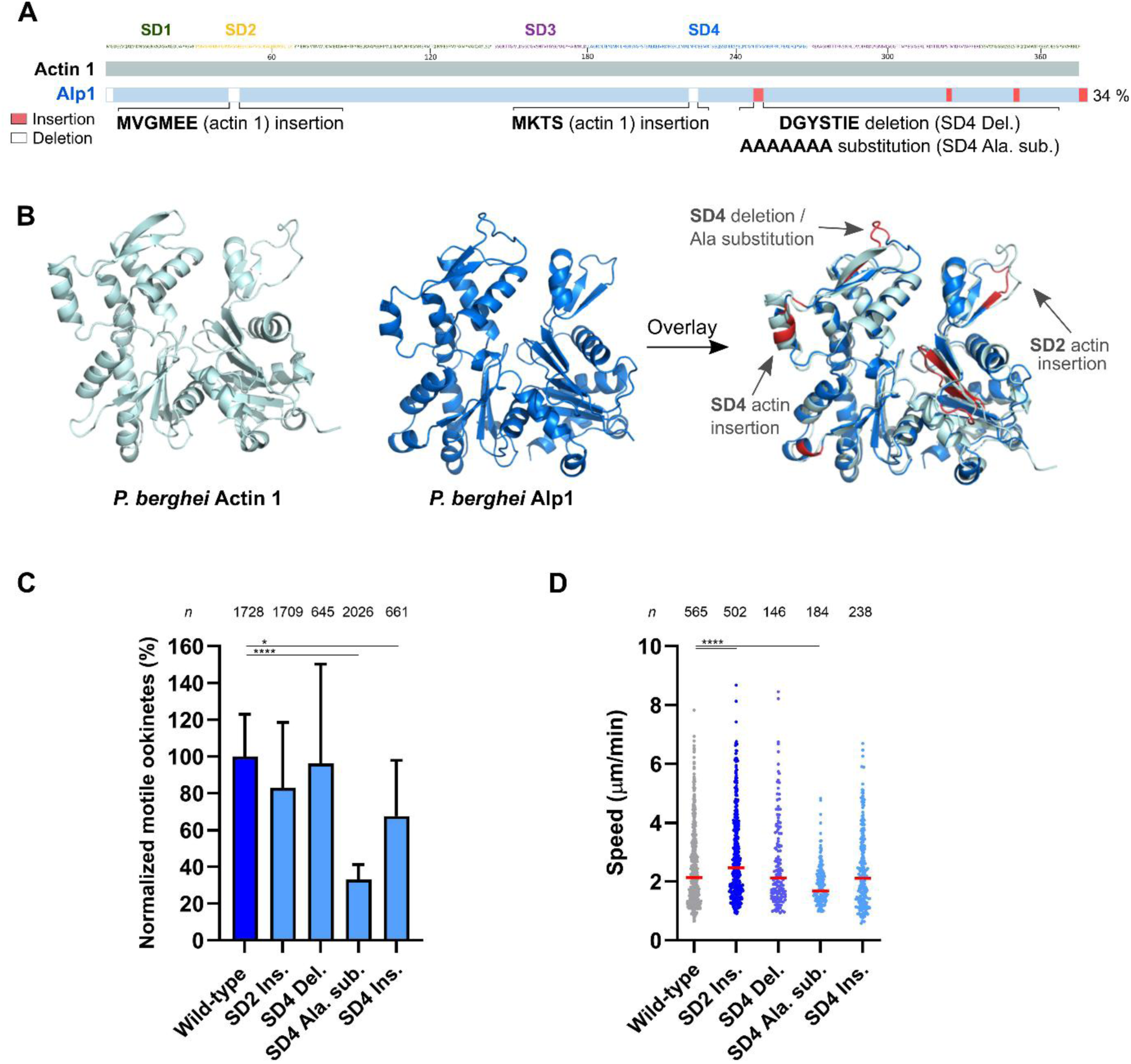
A unique region in subdomain 4 is important for ookinete motility while other regions have minimal impact on Alp1 function. (A) Scheme showing the unique insertions (red boxes) and deletions (white boxes) (InDels) of Alp1 relative to actin 1. The brackets indicate the generated mutations in Alp1 sequences at the corresponding InDel positions. The percentage shows the protein sequence identity between Alp1 and actin1. SD = subdomain. (B) AlphaFold2 structures of Alp1 (dark blue), actin (silver) and overlay of the two. Unique InDel regions (red) are predicted to be generally located within the outer structures of Alp1. Mutations generated are indicated where appropriate. Structures here are the same as in Figure 1. (C) Quantification of motile ookinetes in each Alp1 unique InDel region mutant, normalised to the wild-type motile population. A minor reduction in motile ookinete population was observed for the SD4 Ins. mutant, whereas the motility of the SD4 Ala. sub. mutant was significantly decreased. These results indicate that SD4 of Alp1 contributes to ookinete motility. Fisher’s exact test performed on unnormalized percentage values with each corresponding wild-type control, * p<0.05, **** p<0.0001. (D) Alp1 mutations only modestly affect ookinetes speed. Red line indicates median value. Mann-Whitney test, **** p<0.0001.

To assess the contribution of regions that depart from the classical actin core, regional mutants were generated, whereby Alp1 regions were changed to resemble corresponding actin regions (**Figure S5-S6**). These changes resulted in either an enlarged predicted 1) D-loop (mutant name: SD2 Ins.) or 2) subdomain 4 helix (mutant name: SD4 Ins.). For the 3) insertion region in subdomain 4, we either deleted the region (thereby making it more resemble the actin core, mutant name: SD4 Del.) or, in order to assess the potential loss of interactions of this region while maintaining an insertion, replaced the region with a corresponding amount of alanine residues (mutant name: SD4 Ala. Sub.). Mutations had no impact on the overall predicted Alp1 structure (**Figure S7**). We analyzed the effect of these regional changes on ookinete motility. A very minor effect on ookinete motility for the SD4 Ins. mutant was observed. The alanine substitution of the SD4 insertion region (SD4 Ala. Sub.), but not the deletion of the same insertion region (SD4 Del.), had a striking negative impact on the percentage of motile ookinetes (**Figure 2C-D**), indicating the importance of this region in gliding. None of the mutants phenocopied the knockout, suggesting that a combination of regions or other residues are responsible for Alp1 function.

### The actin chromobody has multiple locations and structures in ookinetes

We thus far identified Alp1 as an essential factor for ookinete motility. Given that actin is a central protein in *Plasmodium* gliding motility^12^ and that many Arps have been shown to interact with actin in classical eukaryotic systems^19,37,38^, we hypothesized that Alp1 was involved in regulation of actin dynamics in ookinetes. In order to observe the alterations of actin in the *alp1* knockout, we employed the actin chromobody, which has previously been used to visualize actin localization in *Toxoplasma gondii*^39,40^, *P. falciparum* blood stages^41^ and *P. berghei* sporozoites^13^. We first assessed the effect and general localization of the probe in ookinetes from wild-type control parasite lines generated previously^13^. Expression of the actin chromobody had no negative effect on ookinete motility (**Figure S8**). We observed diverse sub-cellular localizations of the actin chromobody and classified these on the basis of the position of the highest observed signals. The locations showing the highest proportion of chromobody signal in untreated ookinetes were throughout the whole cell and at the back, while only a small percentage of the population had a “front and back” localization (**Figure 3A-B**). We also observed discrete filamentous structures in the vast majority of untreated ookinetes (**Figure 3A, C**). Treatment with low concentrations of actin modulating compounds altered both signal localization and observed filamentous structures. Treatment with actin filament stabilizing compound jasplakinolide resulted in a shift towards “front and back” localization (**Figure 3A-B**), an observation consistent with *P. berghei* sporozoites^13^ and ookinetes expressing additional copy tagged actin ^42^. We also noticed a change in structures, in which the majority of the treated population had multiple filamentous structures at the back end (67% in 0.5 nM and 80% in 5 nM Jas treated ookinetes), which we refer to as “split filamentous” (**Figure 3A, C**). Treatment with actin filament disruptor cytochalasin D disrupted the filamentous structures and, curiously, rendered an ookinete population that had a diffuse signal with a primary signal collection in the nuclear region (**Figure 3A-C**). Taken together, the actin chromobody is a useful tool for the visualization of actin filaments in ookinetes, revealing primary locations and structures of actin in this stage and can reliably detect shifts in the monomer-filament proportions.

**Figure 3.**
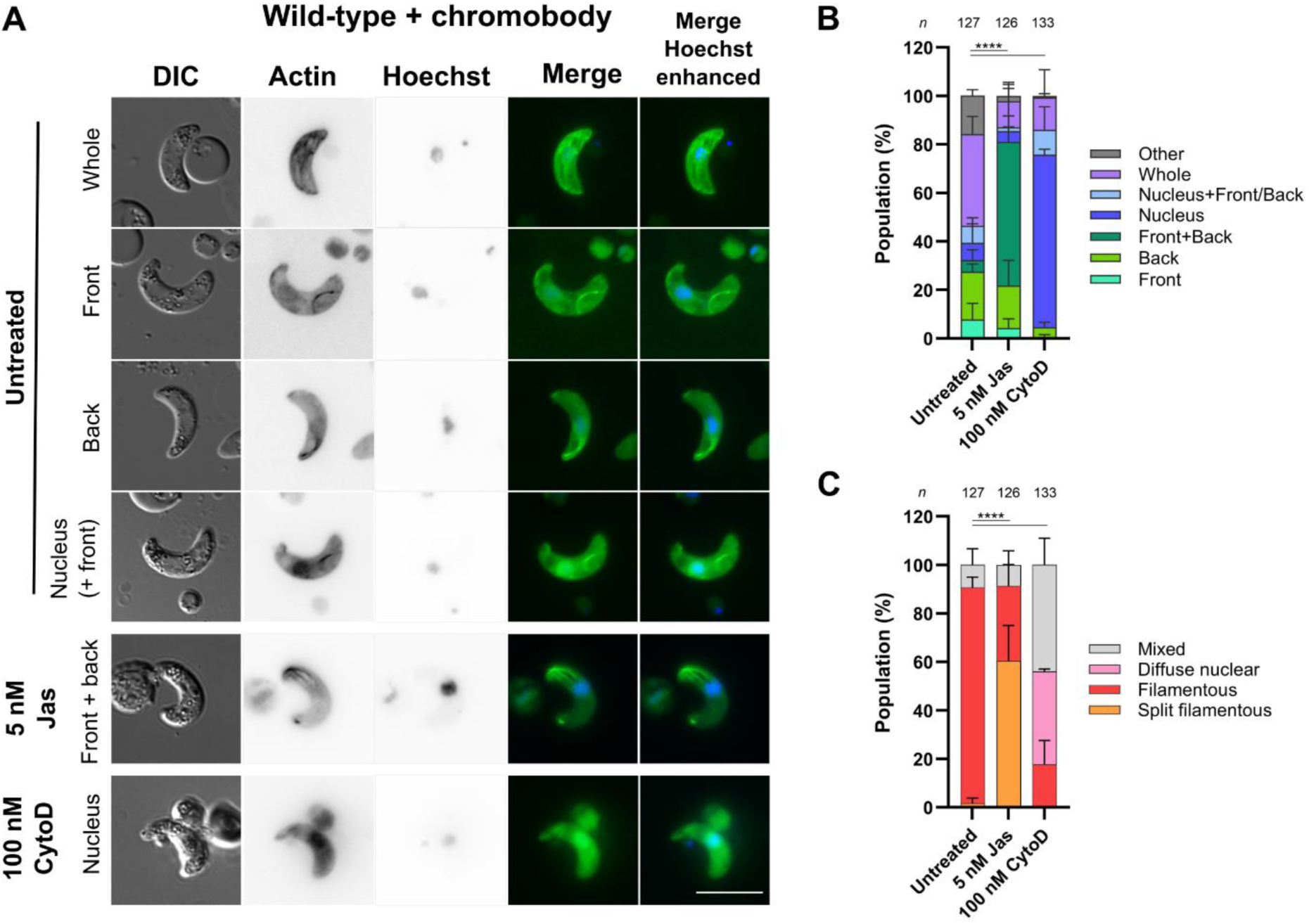
The filament marker actin chromobody shows diverse localisation in ookinetes and is sensitive to actin modulation. (A) Representative fluorescence microscopy images showing the diverse localisation of actin filaments in wild-type ookinetes expressing an actin chromobody. Ookinetes were treated with either 5 nM jasplakinolide (Jas) or 100 nM cytochalasin D (CytoD) or stayed untreated. Scale bar: 10 μm. (B) Quantification of ookinetes based on the location of actin chromobody signal revealed the most common distribution of actin filaments across the entire cell and its posterior region in untreated ookinetes. In contrast, the signals in Jas and CytoD-treated ookinetes were predominantly located at the anterior and posterior regions of the cell (Front+Back) and at the nucleus, respectively. These observations demonstrate the diverse localisation of actin filaments in ookinetes and their sensitivity to classical actin modulators. Fisher’s exact test, **** p<0.0001. (C) Quantification of ookinetes based on the observed structures of actin revealed the presence of filamentous structures in the untreated ookinetes. In Jas-treated ookinetes, split, multi-filamentous signals were predominant, whereas CytoD treatment resulted in a diffuse signal that had increased intensity in proximity to the nucleus. Fisher’s exact test, **** p<0.0001.

### Alp1 potentially facilitates actin filament turnover *in vivo*

We next used the actin chromobody probe to assess changes in actin filament localization and structure in *alp1* knockout parasites. Surprisingly, untreated *alp1* knockout ookinetes resembled the control line in both primary cellular location and the presence of filamentous structures, suggesting that Alp1 does not affect the pattern of actin filament localizations (**Figure 4A-C**). The knockout line also displayed similar structures to controls upon treatment with actin modulating compounds (**Figure 4A**). Noticeably, however, Alp1 knockout ookinetes treated with jasplakinolide showed a significantly higher “front and back” population compared to the corresponding wild-type group. This suggests a more sensitive response to actin stabilization (**Figure 4A-B**). To test sensitivity to modulation, we employed lower concentrations of both modulating compounds and assessed the effect on the observed structures. Treatment with 10 nM cytochalasin D revealed no obvious difference between knockout and control lines. However, treatment with 0.5 nM jasplakinolide resulted in a significantly higher proportion of ookinetes having a split tubular structures compared to the control, which more resembled control with a 10-fold higher jasplakinolide concentration (**Figure 4C**). This combined evidence implies that in Alp1 knockout ookinetes, actin filaments are more prone to stabilization. This suggests that actin filaments might be slightly more stable in the absence of Alp1, hinting to a possible role of this protein in rendering filaments less stable and facilitating filament turnover of actin filaments, which is essential for parasite motility and transmission.

**Figure 4.**
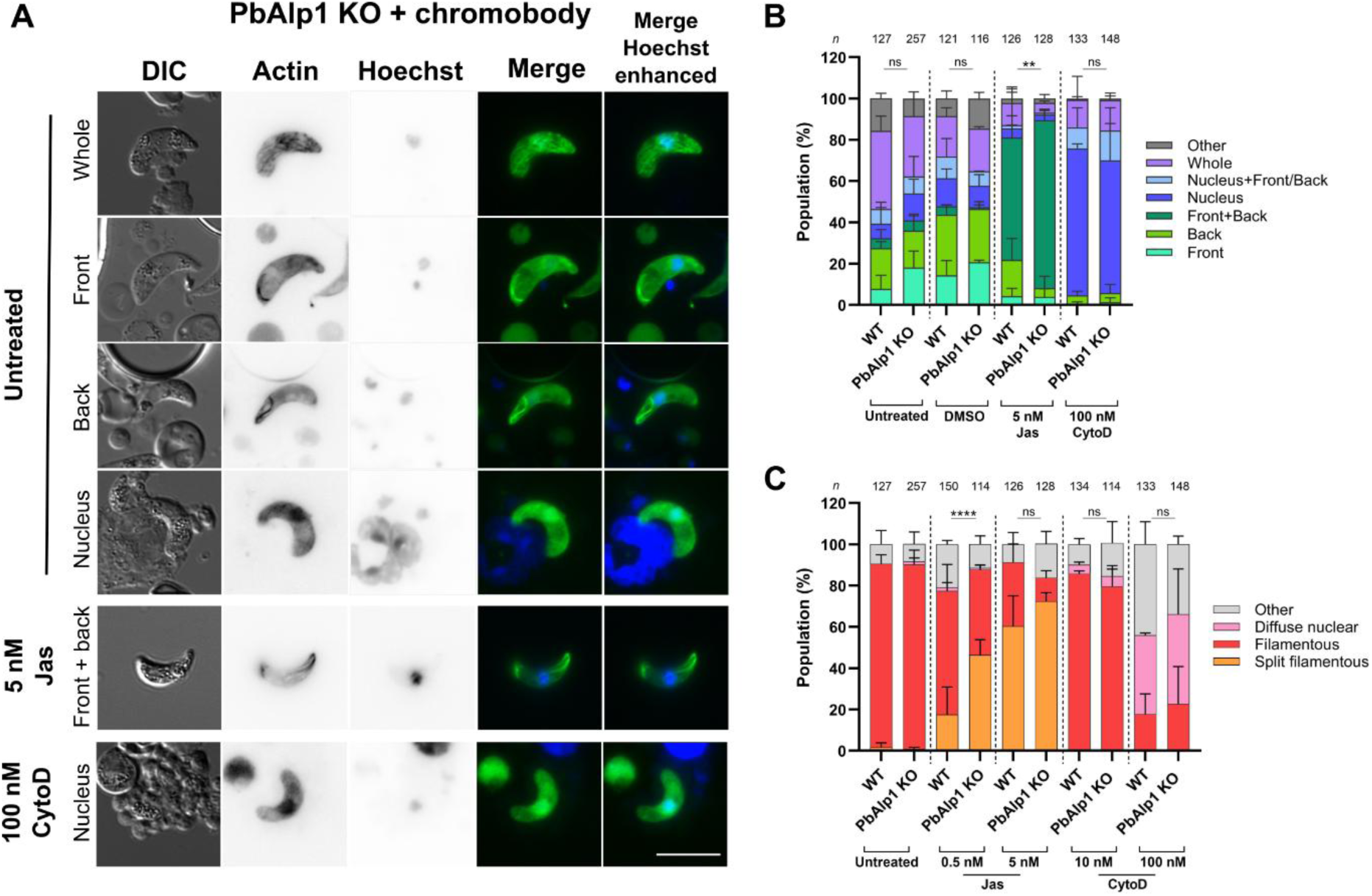
Alp1 deletion renders actin filaments more prone to stabilisation. (A) Representative fluorescence microscopy images showing Alp1 KO ookinetes expressing an actin chromobody, and the effects of treatment with 5 nM Jas and 100 nM CytoD. Scale bar: 10 μm. (B) Comparison of the actin localisation patterns in wild-type and Alp1 KO ookinetes revealed that 5 nM Jas treatment resulted in increased number of front and back signals in the Alp1 KO ookinetes compared to the corresponding wild-type. This indicates that actin localisation in the Alp1 KO ookinetes is more responsive to Jas treatment. Fisher’s exact test, ** p<0.01. Please note that the wild-type untreated, 5 nM Jas and 100 nM CytoD data are the same as from Fig 4B and C. (C) Quantification of the actin filament structures shows that these in the Alp1 KO ookinetes are more susceptible to the filament stabilising effect of Jas, suggesting that Alp1 plays a role in regulating the dynamics of the actin filaments, potentially through filament distabilisation. Fisher’s exact test, **** p<0.0001. Please note that the wild-type untreated, 5 nM Jas and 100 nM CytoD data are the same as from Fig 3B and C.

## Discussion

Using reverse genetics and microscopy, we have discovered an essential role for an apicomplexan-unique actin related protein in malaria transmission. We identified an essential role for Alp1 in ookinete gliding motility while not pronouncedly affecting asexual growth and early mosquito stage parasite development. We further identified a subdomain 4 region on the Alp1 surface that is important for protein function in ookinetes. Finally, we have shown that the actin chromobody is a useful tool to assess actin dynamics in ookinetes and revealed that Alp1 could play a role in actin filament turnover.

### Alp1 and its essential role in parasite progression

Deletion of *alp1* had only a minor effect on blood stage growth and no observable impact on development of early mosquito stage parasites. We generated two different knockout lines, one from the PlasmoGEM resource^33,34^, and the other a complete knockout where all known open reading frame was removed. Phenotypes of both lines were very similar, indicating the robustness of the PlasmoGEM approach in studying actin superfamily members, whereby the actin core is disrupted, and indicating that the remaining C-terminal region is either not sufficiently expressed or not capable of providing any necessary interactions for blood stage growth and gliding. Given the relatively high expression of Alp1 in trophozoites and male gametocytes (**Figure S9**)^31,32^, we were surprised to observe little or no effect at these stages, suggesting potential redundancy with other proteins or a more tolerable molecular requirement at these stages. We further observed the important role of *P. berghei alp1* introns, likely by ensuring sufficient amounts of Alp1 protein needed for ookinete motility. This observation is an important consideration for future design of cross-species complementation constructs.

*Alp1* deletion resulted in a complete block of ookinete motility and thus a block in transmission to mosquitoes, clearly pointing to a role outside of development yet central to parasite motility. These observations are consistent with previous analyses with actin binding proteins and several actin mutants, which showed no measurable effect on blood stages yet strongly affected mosquito stage parasite motility and mosquito organ penetration^12,13,43^. Indeed, a quadruple mutant in actin subdomain 3 also rendered poorly motile ookinetes with reduced oocyst load^13^. These observations point to the fundamental differences in red blood cell invasion and persistent extracellular motility, indicating specialized factors for these processes. For example, merozoites glide at high speed around 1 micron/second for about one minute^44^, ookinetes at a comparatively slow speed of 3-5 microns/minute for over 24 hours^45^ and sporozoites for about 4 hours at 1-2 microns/second^46^. This further highlights the remarkable sensitivity to changes in the gliding machinery and shows the specialized molecular dynamics required for the parasite upon entering the mosquito host. The block in transmission of the *alp1* knockout line meant that we were unable to characterize the role of Alp1 in the sporozoite, a stage that also glides and moves at very high speeds^4,47–49^. Interestingly, *alp1* expression is very low in sporozoites, yet at its highest in liver stages^31,32^ (**Figure S9**). The functional relevance of Alp1 in these stages remains to be determined and could be explored in the future by using promoter-swap or inducible knockout technologies ^50^.

### Comparison with homologues TgAlp1

Two studies of the related *Toxoplasma gondii* Alp1 homologue of Alp1 (TgAlp1) have been conducted^29,30^ showing both similarities and striking differences compared to *Plasmodium*. TgAlp1 was shown to localise in speckle-like structures^30^ and TgAlp1 was further proposed to be part of higher molecular weight complexes^29^. Given that classical Arps typically function in macromolecular assemblies^18,21,51,52^, it is reasonable to suggest that parasite Alps, including Apicomplexan Alp1, could be operating in an analogous manner. Indeed, two other *Plasmodium* Alps were recently shown to operate in a divergent Arp2/3 complex to facilitate nuclear division in activated male gametocytes^28^.

*Tgalp1* was not amenable to deletion using the available technology of the time, while overexpression of TgAlp1 strongly affected *T. gondii* cell division^30^. In contrast, deletion of *P. berghei alp1* did not have an obvious impact on cell division in the parasite stages investigated, but rather on motility of extracellular ookinetes. This might suggest that TgAlp1 has a specialized role in endodyogeny, which is a very different mode of cell division compared to *Plasmodium* schizogony. Whether TgAlp1 also plays a role in *Toxoplasma* tachyzoite gliding motility is currently unknown, and this could now be investigated using for example inducible DiCre approaches^53,54^.

### The subdomain 4 region as a consistent feature important for gliding

Subdomain 4 has also been identified as an important site for actin dynamics. Mutation of *Plasmodium* actin 1 subdomain 4 to corresponding classical actin equivalents influenced motility and invasion of *P. berghei* sporozoites^12^, and it is thus noteworthy that subdomain 4 is emerging as an important site for actin superfamily member interactions in parasite motility. The comparison of predicted 3D structures between actin and Alp1 revealed a very similar structure. However, we also noticed subtle deviations from the canonical actin backbone that we hypothesized were important for Alp1 function. We were therefore surprised to observe that, of the three sites identified and targeted for exchange mutagenesis, only one site pronouncedly affected ookinete motility. This of course does not exclude the importance of other sites in subdomain 2 or 4 and might point to the importance of multiple sites working cooperatively for overall Alp1 function. Indeed, none of the mutations resulted in a phenotype that resembled the knockout lines. We were also surprised to observe a difference between mutations of the one prominent insertion site identified in subdomain 4, where deletion of the insertion did not influence ookinete motility while exchange to a corresponding number of alanines had a striking impact on the proportion of ookinetes able to move productively. The explanation for this is not immediately obvious, but could be due to an alteration on the hydrophobicity profile of the surface of the protein and thus affect the ability of Alp1 to efficiently interact with its required interaction partners^55,56^.

### The actin chromobody is a useful tool in ookinetes and visualizes novel structures that are different between parasite stages

The actin chromobody has been successfully used to visualise actin in different apicomplexans and *Plasmodium* stages^39,13,28^. In *T. gondii*, an elaborate filamentous network was observed in replicating daughter cells^39^. In untreated sporozoites, we previously observed a variety of signal locations with primary collection of chromobody signal at the back of the parasite^13^. In the current study, expression of the chromobody did not negatively influence ookinete motility, indicating no major effect on actin dynamics, and showed striking responsiveness to actin modulating compounds, thus showing that it continues to be a useful tool in visualizing actin filament structures in another parasite stage. We found the presence of long filamentous structures in the majority of parasites, which have thus far not been visualized in ookinetes. Given their thickness, these are presumably bundles of filaments. These structures were not reported in previous work with an anti-actin antibody^42,57^, potentially due to a higher affinity of the chromobody for actin filaments over monomers^39^. Given that ookinetes without these thick filamentous structures were seldom seen, these structures are presumably important for ookinete viability and motility, possibly for transport of key molecules or organelles (**Figure 5**). This also shows the ability for *Plasmodium* actin to be able to form longer filamentous structures and bundles than previously shown *in vitro*^10,58^. Indeed, TIRF microscopy has shown that *Plasmodium* actin 1 can transiently form longer filaments^59^ and more recently, longer filaments and bundles have been observed in untreated *P. falciparum* sporozoites by cryogenic electron tomography^60^. The observed filamentous network is reminiscent of that observed in replicating *T. gondii*, which facilitates the synchronous maturation of daughter cells^39^. This function is not directly relevant for ookinetes as they are a non-replicative stage. However, as with the cytosolic actin network, this network could function in intracellular transport.

**Figure 5.**
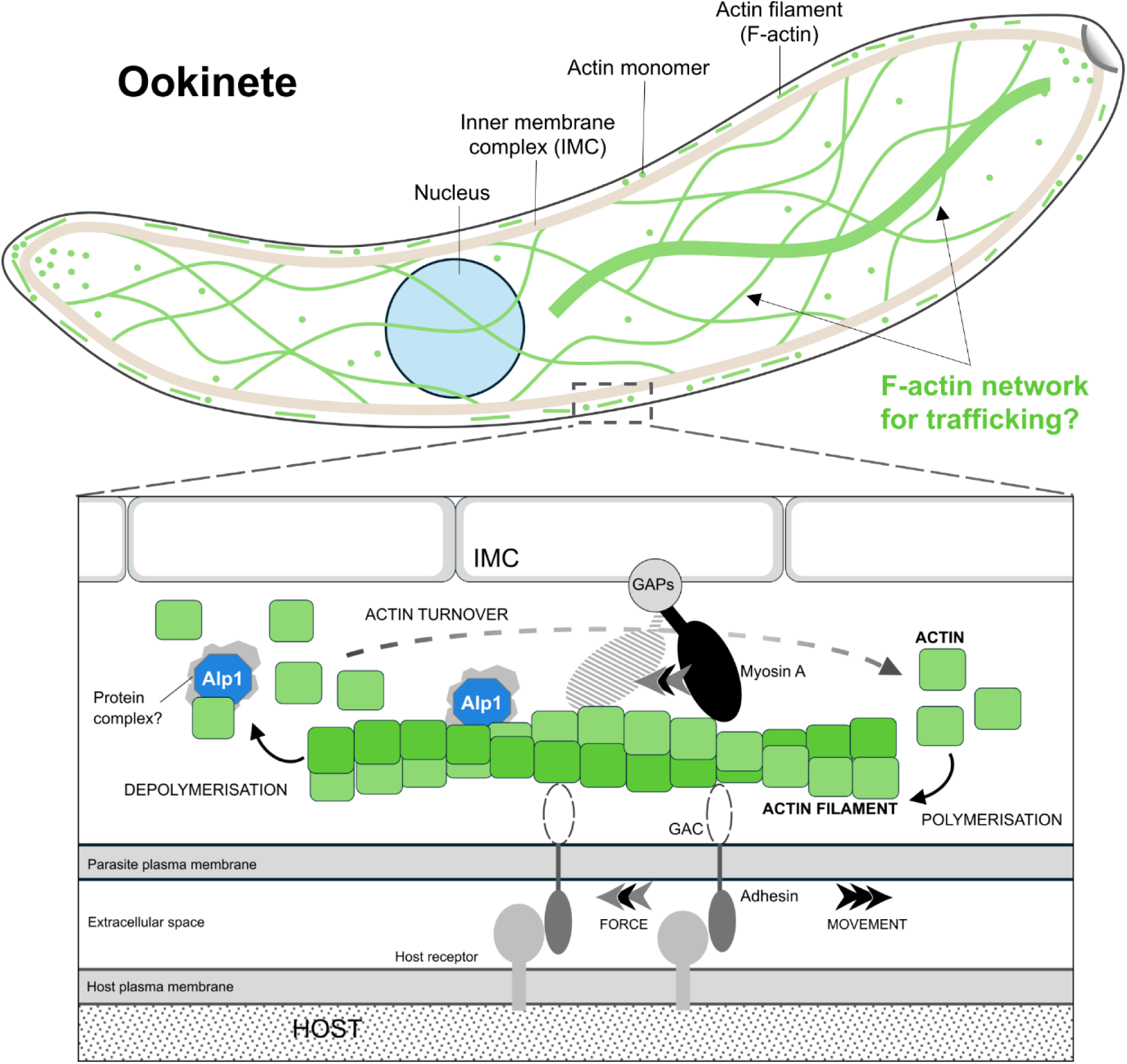
Actin filament dynamics in the ookinete and speculative role of Alp1 in actin turnover. Visualisation of actin using the chromobody revealed an extensive, network-like structure throughout the cell. These structures are likely important for parasite viability and motility and could provide the network for trafficking of important molecules and organelles. We propose that Alp1 (likely in a larger protein complex) supports rapid turnover of actin filaments by destabilising filaments and thus promoting filament disassembly. This may be by directly controlling the rate of dissociation of monomers, interacting with specific actin-binding proteins, or influencing the monomer availability.

Treatment with cytochalasin D resulted in a diffuse localization of the chromobody but with a noticeable signal in the nuclear region. Mammalian stem cells have been shown to have a similar response, whereby treatment with cytochalasin D resulted in disassembly of actin filaments and an increase in transport of actin monomers into the nucleus^61,62^. Interestingly, responsiveness to cytochalasin D was different in sporozoites, which showed an even signal throughout the cell^13^. It is possible, given the importance of actin in chromatin regulation^63^, and the increased DNA content in ookinetes (which have 4 copies of the genome), that ookinetes may have a more efficient actin import system compared to sporozoites (which are haploid). Treatment with filament stabilizing compound jasplakinolide resulted in a shift in signal locations (to front and back) and structure (towards a split filamentous arrangement). This shift in location is consistent with that observed in sporozoites^12,13^. A split arrangement was sometimes observed in jasplakinolide treated sporozoites expressing an additional copy of actin^12^ and likely reflects the accumulation of actin in the primary sites of actin assembly and disassembly. Interestingly, a dot-like structure observed here in ookinetes was also observed in the front of sporozoites^13^. A formin isotype has been localized at the apical end of gliding sporozoites^12^ and in free merozoites^64^. While formins have not yet been localized in ookinetes, this suggests the presence of an actin nucleator at the front of ookinetes and an overall similar sub-cellular arrangement in the gliding machinery^60,65^.

The use of the chromobody allows for comparison of two different stages with two different speeds and persistence requirements. While ookinetes displayed observable actin structures, sporozoites typically displayed more “blob”-like localizations indicative of “pools” of actin filaments^13^. Nonetheless, both stages had the back as one of the major sites of F-actin signal and a discrete point on the apex with jasplakinolide treatment. Taken together, the actin chromobody observations begin to point out the similarities and differences in these two motile stages: Both make use of similar locations for performing motility. One the other hand, the presence of filamentous structures in the ookinetes points to a likely more static cytoskeleton in ookinetes in comparison to sporozoites. As discussed above, these structures are likely a requirement for ookinete viability and motility, but could also additionally reflect the extent of rapid actin dynamics between the stages. Sporozoites glide more than an order of magnitude faster than ookinetes^48,49,66^. Thus, the sporozoite would require much more actin turnover to move as fast as possible and probably would not allow the formation of more obvious, thicker filamentous structures. In contrast, ookinetes move slower in the soft environment of the blood bolus^67^ and much longer^45^ and thus could have less demands for very rapid actin turnover. This is consistent with our previous work with mutations in actin, whereby most mutations that were viable did not affect ookinete motility and infection, but pronouncedly affected sporozoite motility^12,13^. Nonetheless, our work emphasizes the need for ookinetes to have efficient and coordinated actin turnover, even with relatively slower speeds. In line with this, we previously observed a quadruple actin mutant which was not able to efficiently glide at the ookinete stage^13^, indicating a specialist actin dynamic required for gliding.

### The role of Alp1 in ookinete motility

Assessing the chromobody in the Alp1 knockout background indicated a potential role of Alp1 in regulating ookinete actin dynamics. Our data points to a subtle yet critical role in actin regulation. Since there was a noticeable response of actin filaments to low concentrations of jasplakinolide in the *alp1* knockout line, it appears that actin filaments *in vivo* are more prone to being stable in the absence of Alp1. Consequently, we propose that Alp1 might be involved in mediating filament turnover and thus, directly or indirectly, be a type of filament depolymerizer. Indeed, yeast Arp4, working in concert with Arp8, has been reported to depolymerize actin filaments^38^. Curiously, Arp4 also has an insertion site in the subdomain 4 region^38^, suggesting a possible conserved structural role of this site in actin depolymerization. We speculate that, given that many Arps have been documented to interact with actin, that this could be a direct interaction (**Figure 5**) like in the case of Arp4, but detailed biochemical investigations would be required to test this hypothesis.

Taken together, our work has identified an essential apicomplexan unique Arp that has evolved specialist features to facilitate ookinete gliding motility through regulating actin dynamics and malaria transmission.

## Methods

### Animal experiments

All animal experiments were performed according to the Federation of European Laboratory Animal Science Associations (FELASA) and Gesellschaft für Versuchstierkunde/Society for Laboratory Animal Science (GV-SOLAS) standard guidelines. Approval was granted by the responsible German authorities (Ethics Committees: Regierungsprasidium Giessen and Regierungsprasidium Karlsruhe). For all experiments, female four to six week old Swiss mice were used (obtained from Janvier Laboratories, France). *Anopheles stephensi* mosquitoes were reared and maintained by standard breeding methods.

### Generation of PbAlp1 knockout, complementation and mutant lines

The pGEM KO construct for *P. berghei* Alp1 (PBANKA_0936900) was obtained from PlasmoGEM (https://plasmogem.umu.se/pbgem/)^33,34^. For the complete removal of PbAlp1, the 5’ and 3’ untranslated regions (UTRs) flanking *alp1* were PCR amplified from genomic DNA (gDNA) using the appropriate primers (see Table S1 for a full list of primers used). These were then cloned into the Pb262 transfection vector ^68^ using NotI/EcoRV (5’ UTR) and ApaI/PmeI (3’ UTR) restriction sites (all enzymes purchased from New England Biolabs, Germany). Consequently, the transfection construct contained the *hdhfr-yfcu* selection cassette^69^, which was flanked by the 5’ and 3’ UTRs of *alp1*. The construct was linearised using NotI and PmeI, ethanol-precipitated, and then transfected into the recipient wild-type *P. berghei* ANKA strain, as previously described ^70^. Positive selection was achieved using 0.07 mg/mL pyrimethamine. Correctly integrated recipient line was isolated by limiting dilution and subsequent genotyping PCR. All the constructs in the following sections were generated using the same techniques for transfection, selection and limiting dilution. To remove the selection cassette, selected recipient parasite line was selected using 1 mg/mL 5-fluorocytosine, followed by limiting dilution and genotyping PCR. Asexual growth rates were calculated as described previously during limiting dilution^71^, based on the parasitaemia obtained one day before reaching 1%.

*P. berghei* Alp1 complementation was achieved using the Pb238 vector ^12,13,72,73^, which carries the *alp1* open reading frame (ORF) and its flanking 5’ and 3’ UTRs, as well as the *hdhfr* selection cassette downstream. The 5’ UTR-*alp1*, 3’ UTR and additional flanking 3’ UTR fragments were PCR amplified from gDNA and cloned into the transfection vector using the SacII/BamHI (5’ UTR-*alp1*), BamHI/EcoRV (3’ UTR) and AvrII/KpnI (flanking 3’ UTR) restriction sites. The construct was linearised using SacII and KpnI enzymes prior to transfection.

*P. falciparum alp1* was amplified and complemented as complementary DNA (cDNA) to exclude the exons. The target gene was expressed using the endogenous *pbalp1* 5’ and 3’ UTRs, which were amplified from the respective gDNA. The amplified fragments were then integrated into the Pb262 transfection vector via the following restriction sites: BssHII/NotI (5’ UTR), NotI/EcoRI (*pfalp1*), EcoRI/EcoRV (3’ UTR) and FseI/PmeI (flanking 3’ UTR). The resulting construct contained *pfalp1* ORF flanked by *pbalp1* 5’ and 3’ UTRs, the *hdhfr-yfcu* selection cassette^69^ and an additional 3’ UTR to enclose the construct. BssHII and PmeI enzymes were used to linearise the transfection construct.

PfEx.+PbInt. Alp1 complementation construct was generated using the overlapping PCR technique. *P. falciparum alp1* exons and *P. berghei alp1 introns* were amplified from the corresponding gDNA with primer sets bearing overlapping sequences with neighbouring fragments. The Pf-Pb hybrid *alp1* frangment was cloned into the Pb238 transfection vector, which already contained the 5’ and 3’ *pbalp1* UTRs from the previous Alp1 complementation study. This was done using the BssHII and BglII restriction sites.

Similarly, all Alp1 unique region mutants were generated via overlapping PCR by amplifying gDNA with primer sets containing target mutations in the overlapping sequences. The *alp1* mutant fragments were then inserted between the BssHII/BamHI restriction sites of the aforementioned Pb238 complementation transfection vector. Both the PfEx.+PbInt. and the unique region mutant Alp1 constructs were linearised using the BssHI and PmeI enzymes before transfection. All linearised Alp1 complementation and mutant constructs were transfected into recipient Alp1 full KO *P. berghei* ANKA strain and subsequently selected with pyrimethamine as above.

### Generation of PbAlp1 knockout chromobody line

For the generation of the Alp1 KO parasite line expressing the actin chromobody, the Pb238 actin chromobody transfection vector^13^ was linearised using the SalI and SapI restriction sites, then transfected into the chromosome 12 of the recipient Alp1 full KO *P. berghei* ANKA strain. This was followed by positive selection using pyrimethamine, as described in a previous section. The control actin chromobody-expressing wild-type *P. berghei* ANKA parasite line used in this study was generated and provided by Yee et al. (2022).

### Mosquito infection

Prior to mosquito feeding, a cryostock of the required parasite line was injected i.p. into a mouse. Four days post-infection, the infected donor mouse was bled via cardiac puncture and 20x10^6^ blood stage parasites transferred into two naïve recipient mice through i.p. injection. Three days post-transfer, exflagellation was assessed by incubation of a drop of tail blood for 10 minutes at 20°C, in the dark. Exflagellation events were observed by light microscopy (Carl Zeiss GmBH, Germany). If two or more exflagellation events per field were observed, a mosquito cage feed was performed. Adult mosquitos aged between between 3 to 7 days post-hatching were starved for at least 6 hours to increase feeding efficiency. Mice were anesthetised with ketamine/xylazine by i.p. injection (100 mg/kg ketamine, 3 mg/kg xylazine). Completely anesthetised mice were then placed on top of the mosquito cage and mosquitos were allowed to feed for up to 30 minutes. To allow feeding of all mosquitos, the mice position on the cage was changed every 10 minutes. Once infected, mosquitos were housed at 21 °C and 70 % humidity, fed with pads saturated with 10% (w/v) saccharose with 0.05% PABA and 1% (w/v) NaCl.

### Quantification of oocyst formation

To check whether parasites are capable of forming oocysts, midguts of mosquitos were dissected between day 10 and 14 post mosquito feeding. Midguts were dissected into 100 μl of PBS on ice and subsequently permeabilised with 1 % (v/v) Nonidet P40 for 20 minutes at room temperature. This was followed by staining with 0.1 % (w/v) mercurochrome for 30-120 minutes at room temperature. After staining, midguts were washed with PBS until the supernatant became clear and subsequently placed onto a glass slide. Stained midguts were imaged with the wide-field microscope (Carl Zeiss, Axiovert 200M, Germany) at 10x magnification using a green filter (38 HE Green Fluorescent Prot). The number of oocysts per infected midgut and the infected midguts of total midguts were quantified.

### Ookinete culture and motility assay

Ookinete cultures were performed similar to that described previously^13^. Ookinetes were developed directly from whole blood, whereby a donor mouse was infected by i.p. injection of a cryo-preserved parasite line. Parasites were allowed to grow in the infected mouse until a parasitaemia of 1.5% to 2%. The parasites were harvested from the infected mouse via cardiac puncture and the blood was used for fresh blood transfer of 20x10^6^ blood stage parasites intraperitoneally into naïve recipient mouse. Three days post-transfer, exflagellation centres were assessed. When more than one exflagellation event per field was observed, the parasites were harvested from the recipient mouse via cardiac puncture and directly transferred to 12 mL pre-incubated ookinete medium (250 mL RPMI1640 + HEPES + glutamine, 12.5 mg hypoxanthine, 2.5 ml penicillin/streptomycin (100x), 0.5 g sodium bicarbonate, 5.12 mg, xanthurenic acid (100 μM), 16% (v/v) foetal bovine serum, pH 7.8) at 19°C. The parasites were cultured at 19°C for 21 hours. Following the incubation, 1 mL aliquot of the ookinete culture was taken and pelleted at 7000 rpm for 2 minutes at room temperature. The supernatant was removed and 3 μL of the pellet was used to make a Giemsa-stained smear to check for ookinete development.

To set up an ookinete motility assay, Nycodenz purification was performed. 10 mL of the ookinete culture was carefully underlaid with 10 mL of prewarmed (19°C) 63% (v/v) Nycodenz cushion in a 50 mL tube and spun at 1000 rpm for 25 minutes at room temperature (no brake). After centrifugation, ookinetes enriched at the interphase were harvested using a Pasteur pipette into a 15 mL tube. The purified ookinetes were pelleted at 1 000 rpm for 8 minutes at room temperature. The supernatant was removed and the pellet was resuspended with 1 mL of ookinete medium. During the assay, purified ookinetes were kept at 19°C in the dark.

For imaging motile ookinetes, a 200 μL aliquot was briefly centrifuged for 10 seconds at 13400 rpm, 190 µL of the supernatant was removed and the remaining 5 -10 µL were used to gently resuspend the ookinete pellet. Approximately 2 μL of this sample was then mounted on a microscope slide, covered with a cover slip and sealed with VALAP (10 g Vaseline, 10 g paraffin, 10 g lanolin). Ookinete motility was visualised using a wide-field microscope (Carl Zeiss Axio Observer Z1 with or Axioskop 2 MOT) with 40x objective. Images were acquired every 30 seconds for 15 minutes in the DIC channel (50 ms exposure). Two 15-minute movies were made from each aliquot and up to five technical repeats (i.e. 5x200 µl preparations) for each purification were performed. At least two biological repeats were performed for each line. Motile ookinetes were defined as those ookinetes that moved more than one parasite length. Average speed of ookinetes were calculated using the Fiji Manual Tracking plug-in ^74^.

### Fluorescence microscopy

Prior to immunostaining, the purified ookinetes were pelleted and fixed in a 4% (w/v) paraformaldehyde and 0.05% (v/v) 25% glutardialdehyde solution for at least 60 minutes at room temperature. The fixed ookinetes were then washed three times in PBS (centrifugation: 7000 rpm for 2 minutes at room temperature; the same conditions apply to all subsequent centrifugation steps unless otherwise specified) and stored in fresh PBS at 4 °C.

Both wild-type and Alp1 KO actin chromobody ookinetes were purified and fixed as previously described. For the untreated sample, 100–300 µL of purified ookinetes were centrifuged at maximum speed for 20 seconds and resuspended in 500 µL PFA fixing solution. For jasplakinolide and cytochalasin D treatment, 198 μL of purified ookinetes were supplemented with 2 μL of 50 nM or 500 nM jasplakinolide in DMSO (for the final concentration of 0.5 nM and 5 nM, respectively) or 2 μL of 1 μM or 10 μM cytochalasin D in DMSO (for a final concentration of 10 nM and 100 nM, respectively). After a 3-minute incubation with the compounds, the samples were centrifuged for 20 seconds at maximum speed and resuspended in 500 μL PFA fixing solution. All ookinete samples were fixed for a minimum of 1 hour at room temperature and washed three times with 1 mL PBS. The samples were then resuspended in 100 μL PBS supplemented with 5 μg/mL Hoechst dye. After incubating for 20 minutes at room temperature, the sample was washed once more with 1 mL of PBS, followed by removal of the supernatant leaving approximately 10–30 μL of solution with the pellet. For imaging, 2 μL of the pellet was mounted onto a glass slide with a cover slip. Fluorescence images were acquired using an Axio Observer Z1 microscope with a 63x oil objective. The actin-chromobody and Hoechst signals were imaged using the EGFP (150 ms exposure time) and H3258 (25 ms exposure time) filters, respectively. The Fiji ImageJ software (Schneider, C. A. et al., 2012) was employed to extract fluorescent ookinete images from the multi-channel raw microscopy data.

### Prediction and manipulation of protein structures

Predicted structures of *Plasmodium* Alp1 and the mutants were generated using the AlphaFold2 online tool^75^. To generate predictions for the mutated versions of PbAlp1, the corresponding amino acid sequences were provided to the programme. The protein model with the highest prediction confidence was selected for use in the experimental planning and presentation. The protein structures were coloured, highlighted and overlaid using open source PyMOL.

## Acknowledgments

This study was funded by the LOEWE Centre DRUID (“Novel Drugs Targets against Poverty-related and Neglected Tropical Infectious Diseases”) within the Hessian Excellence Program. We thank J. Niermann, T. Holzapfel and E. Nikolaou for technical assistance. We also thank J. Przyborski and C. Grevelding for helpful discussions during the course of the study.

## Supplementary figures and tables

**Table S1: Primers used in the study**

**Figure S1:** Generation and genotyping of Alp1 KO lines.

(A) Integration schemes and the genomic locus arrangement in both pre-transfected and transfected lines of PbAlp1 KO-PG. The orientation of the small arrows is indicative of the promoter, while the numbers and the dotted lines indicate the primer pairs used (see **Table S1**) and their coverage for genotyping. (B) Representative agarose gels of the selected genotyped parasite lines. Schemes below illustrate the structures of endogenous and the partially deleted *alp1*. (C) Integration schemes and the genomic locus arrangement in both pre-transfected and transfected lines of PbAlp1 KO-full. The orientation of the small arrows is indicative of the promoter, while the numbers and the dotted lines indicate the primer pairs used (see **Table S1**) and their coverage for genotyping. (D) Representative agarose gels of the selected genotyped parasite lines.

**Figure S2:** Generation and genotyping of PbAlp1 complementation line.

(A) Integration schemes and the genomic locus arrangement in both pre-transfected (PbAlp1 KO-PG) and transfected lines of PbAlp1 complementation. The orientation of the small arrows is indicative of the promoter, while the numbers and the dotted lines indicate the primer pairs used (see **Table S1**) and their coverage for genotyping. (B) Representative agarose gels of the selected genotyped parasite lines.

**Figure S3:** Generation and genotyping of PfAlp1 cDNA complementation line.

(A) Integration schemes and the genomic locus arrangement in both pre-transfected (PbAlp1 KO-full) and transfected lines of PfAlp1 cDNA complementation. The orientation of the small arrows is indicative of the promoter, while the numbers and the dotted lines indicate the primer pairs used (see **Table S1**) and their coverage for genotyping. (B) Representative agarose gels of the selected genotyped parasite lines.

**Figure S4:** Generation and genotyping of PfEx.+PbIn. Alp1 complementation line.

(A) Integration schemes and the genomic locus arrangement in both pre-transfected (PbAlp1 KO-full) and transfected lines of PfEx.+PbIn. Alp1 complementation. The orientation of the small arrows is indicative of the promoter, while the numbers and the dotted lines indicate the primer pairs used (see **Table S1**) and their coverage for genotyping. (B) Representative agarose gels of the selected genotyped parasite lines.

**Figure S5:** Sequence alignments of PbAlp1 with actin and Alp1 from other *Plasmodium* species.

(A) Protein sequence alignment of *P. berghei* actin 1 and Alp1, with the mutated insertion and deletion (InDel) regions indicated. For the actin insertion mutants (indicated in blue), amino acid residues from actin 1 were inserted into the Alp1 sequence: MVGMEE in the D-loop and MKTS in subdomain 4. Two mutations were introduced in the second deletion site in subdomain 4 (indicated in red): In one mutant, the amino acid residues in the Alp1 sequence (DGYSTIE) were deleted, while in the other, these residues were substituted with alanine. D-loop = DNase-I binding loop; SD = subdomain. (B) Predicted structure of PbAlp1 with InDel sites compared to Pbactin 1. Structure prediction was performed using AlphaFold2 (Jumper et al. 2021). (C) Protein sequence of Pbactin 1 aligned with Alp1 sequences from different *Plasmodium* species. Several unique insertions (red frame) and deletions (blue frames) are present in Alp1 in different *Plasmodium* species, compared to actin 1 in *P. berghei*. These InDels are highly conserved across species. The top row shows Pbactin 1, followed by Alp1 sequences from other *Plasmodium* species. All sequence alignments were conducted using Clustal Omega ^76^.

**Figure S6:** Generation and genotyping of Alp1 InDel mutant lines.

(A) Integration schemes and the genomic locus arrangement in both pre-transfected (PbAlp1 KO-full) and transfected lines of Alp1 InDel mutants. The orientation of the small arrows is indicative of the promoter, while the numbers and the dotted lines indicate the primer pairs used (see **Table S1**) and their coverage for genotyping. (B) Representative agarose gels of the selected genotyped parasite lines from each InDel mutant.

**Figure S7:** Predicted structural effects of mutations in Alp1.

In the top row, the enlarged images of the predicted Alp1 mutation sites taken from alignment of Pbactin 1 (green) and PbAlp1 (light blue) (**Figure S8**) are shown. Insertions are marked in dark blue or red as indicated by the description. The two middle panels with the PbAlp1 subdomain 4 (SD4) insertion are identical as two different mutations were generated at this site. In the bottom row, the mutated Alp1 sites (grey) are aligned with the predicted wild type (light blue) structures. The actin insertions are marked in blue and the alanine substitution in orange. All structure predictions were performed with AlphaFold2 and alignments were performed with open source PyMOL. As the top and bottom pictures were positioned and enlarged manually, minor differences in the size and tilt of the depicted protein regions could not be prevented. The mutations had no major effects on the predicted overall structures.

**Figure S8:** Impact of the actin chromobody on ookinete motility and generation and genotyping of the Alp1 KO actin chromobody line.

(A, B) Quantification of ookinete motility and speeds in the “wild-type” (i.e. no changes made in other loci) expressing actin chromobody line indicated that the chromobody has no negative effect on ookinete motility. The dotted line indicates the wild-type (no chromobody) data as in Figure 2B-C. Statistical analysis represents a level of significance against wild-type motile ookinetes and speeds (in A, Fisher’s exact test, * p<0.05; in B, Mann-Whitney test for the speed, **p<0.01). (C) Integration schemes and the genomic locus arrangement in both pre-transfected and transfected lines of Alp1KO-actin chromobody. The orientation of the small arrows is indicative of the promoter, while the numbers and the dotted lines indicate the primer pairs used (see **Table S1**) and their coverage for genotyping. (D) Representative agarose gels of the selected genotyped parasite lines. To ensure the absence of Alp1 in the transfected line, the corresponding locus was additionally genotyped.

**Figure S9:** RNASeq profile of *P. berghei alp1* (PBANKA_0936900) throughout the life cycle. The bar graph, taken from the SPOT analysis (https://frischknechtlab.shinyapps.io/SPOT/) (Farr et al., 2021; Howick et al., 2019), shows that the *alp1* gene is highly expressed at the liver, trophozoite and male gamete stages. Meanwhile, it is less expressed in highly motile ookinetes and sporozoites.

## Notes

### Competing Interest Statement

The authors have declared no competing interest.

## References

1. Smith, M. L. & Styczynski, M. P. Systems Biology-Based Investigation of Host-Plasmodium Interactions. Trends Parasitol 34, 617–632 (2018).

2. Kuehn, A. & Pradel, G. The coming-out of malaria gametocytes. J Biomed Biotechnol 2010, 976827 (2010).

3. Patra, K. P. et al. A Hetero-Multimeric Chitinase-Containing Plasmodium falciparum and Plasmodium gallinaceum Ookinete-Secreted Protein Complex Involved in Mosquito Midgut Invasion. Front Cell Infect Microbiol 10, 615343 (2021).

4. Frischknecht, F. & Matuschewski, K. Plasmodium Sporozoite Biology. Cold Spring Harb Perspect Med 7, a025478 (2017).

5. Douglas, R. G., Amino, R., Sinnis, P. & Frischknecht, F. Active migration and passive transport of malaria parasites. Trends Parasitol 31, 357–62 (2015).

6. Heinzelman, M. B. Gliding motility in apicomplexan parasites | Elsevier Enhanced Reader. Semin. Cell Dev. Biol. 46, 135–142 (2015).

7. Singer, M. & Frischknecht, F. Still running fast: *Plasmodium* ookinetes and sporozoites 125 years after their discovery. Trends in Parasitology 39, 991–995 (2023).

8. Douglas, R. G., Moon, R. W. & Frischknecht, F. Cytoskeleton Organization in Formation and Motility of Apicomplexan Parasites. Annu Rev Microbiol. 78, 311–335 (2024).

9. Kobayashi, Y. & Douglas, R. G. Highly divergent apicomplexan cytoskeletons provide additional models for actin biology. FEBS J 10.1111/febs.70263 (2025) doi:10.1111/febs.70263.

10. Vahokoski, J. et al. Structural differences explain diverse functions of Plasmodium actins. PLoS Pathog 10, e1004091 (2014).

11. Schüler, H., Mueller, A.-K. & Matuschewski, K. Unusual properties of Plasmodium falciparum actin: new insights into microfilament dynamics of apicomplexan parasites. FEBS Lett 579, 655–660 (2005).

12. Douglas, R. G. et al. Inter-subunit interactions drive divergent dynamics in mammalian and Plasmodium actin filaments. PLoS Biol. 16, e2005345 (2018).

13. Yee, M., Walther, T., Frischknecht, F. & Douglas, R. G. Divergent Plasmodium actin residues are essential for filament localization, mosquito salivary gland invasion and malaria transmission. PLoS Pathog 18, e1010779 (2022).

14. Sattler, J. M., Ganter, M., Hliscs, M., Matuschewski, K. & Schüler, H. Actin regulation in the malaria parasite. European Journal of Cell Biology 90, 966–971 (2011).

15. Muller, J. et al. Sequence and comparative genomic analysis of actin-related proteins. Mol. Biol. Cell 16, 5736–48 (2005).

16. Schafer, D. A. & Schroer, T. A. Actin-related proteins. Annu. Rev. Cell Dev. Biol. 15, 341–63 (1999).

17. Hammesfahr, B. & Kollmar, M. Evolution of the eukaryotic dynactin complex, the activator of cytoplasmic dynein. BMC Evol Biol 12, 95 (2012).

18. Reck-Peterson, S. L., Redwine, W. B., Vale, R. D. & Carter, A. P. The cytoplasmic dynein transport machinery and its many cargoes. Nat Rev Mol Cell Biol 19, 382–398 (2018).

19. Urnavicius, L. et al. The structure of the dynactin complex and its interaction with dynein. Science 347, 1441–1446 (2015).

20. Francis, J. et al. Activation of Arp2/3 complex by a SPIN90 dimer in linear actin-filament nucleation. Nat Struct Mol Biol 1–13 (2025) doi:10.1038/s41594-025-01673-8.

21. Gautreau, A. M., Fregoso, F. E., Simanov, G. & Dominguez, R. Nucleation, Stabilization and Disassembly of Branched Actin Networks. Trends Cell Biol 32, 421–432 (2022).

22. Liu, T. et al. Arp2/3-mediated bidirectional actin assembly by SPIN90 dimers. Nat Struct Mol Biol 1–10 (2025) doi:10.1038/s41594-025-01665-8.

23. Mullins, R. D., Heuser, J. A. & Pollard, T. D. The interaction of Arp2/3 complex with actin: Nucleation, high affinity pointed end capping, and formation of branching networks of filaments. Proc. Natl. Acad. Sci. U.S.A. 95, 6181–6186 (1998).

24. Oma, Y. & Harata, M. Actin-related proteins localized in the nucleus: from discovery to novel roles in nuclear organization. nucleus 2, 38–46 (2011).

25. Sen, B. et al. Nuclear actin structure regulates chromatin accessibility. Nat Commun 15, 4095 (2024).

26. Shen, X., Mizuguchi, G., Hamiche, A. & Wu, C. A chromatin remodelling complex involved in transcription and DNA processing. Nature 406, 541–544 (2000).

27. Gordon, J. L. & Sibley, L. D. Comparative genome analysis reveals a conserved family of actin-like proteins in apicomplexan parasites. BMC Genomics 6, 179 (2005).

28. Hentzschel, F. et al. An atypical Arp2/3 complex is required for Plasmodium DNA segregation and malaria transmission. Nat Microbiol 10, 1775–1790 (2025).

29. Gordon, J. L., Buguliskis, J. S., Buske, P. J. & Sibley, L. D. Actin-like protein 1 (ALP1) is a component of dynamic, high molecular weight complexes in Toxoplasma gondii. Cytoskeleton 67, 23–31 (2010).

30. Gordon, J. L., Beatty, W. L. & Sibley, L. D. A novel actin-related protein is associated with daughter cell formation in Toxoplasma gondii. Eukaryot Cell 7, 1500–12 (2008).

31. Howick, V. M. et al. The Malaria Cell Atlas: Single parasite transcriptomes across the complete Plasmodium life cycle. Science 365, (2019).

32. Farr, E. B., Sattler, J. M. & Frischknecht, F. SPOT: a web-tool enabling swift profiling of transcriptomes. Bioinformatics 38, 284–285 (2021).

33. Bushell, E. et al. Functional Profiling of a Plasmodium Genome Reveals an Abundance of Essential Genes. Cell 170, 260–272.e8 (2017).

34. Pfander, C. et al. A scalable pipeline for highly effective genetic modification of a malaria parasite. Nat Methods 8, 1078–1082 (2011).

35. Spaccapelo, R. et al. Plasmepsin 4-deficient Plasmodium berghei are virulence attenuated and induce protective immunity against experimental malaria. Am. J. Pathol. 176, 205–17 (2010).

36. Bennink, S., Kiesow, M. J. & Pradel, G. The development of malaria parasites in the mosquito midgut. Cell. Microbiol. 18, 905–918 (2016).

37. Rotty, J. D., Wu, C. & Bear, J. E. New insights into the regulation and cellular functions of the ARP2/3 complex. Nat Rev Mol Cell Biol 14, 7–12 (2013).

38. Fenn, S. et al. Structural biochemistry of nuclear actin-related proteins 4 and 8 reveals their interaction with actin. EMBO J. 30, 2153–2166 (2011).

39. Periz, J. et al. Toxoplasma gondii F-actin forms an extensive filamentous network required for material exchange and parasite maturation. eLife 6, e24119 (2017).

40. Periz, J. et al. A highly dynamic F-actin network regulates transport and recycling of micronemes in Toxoplasma gondii vacuoles. Nat Commun 10, 4183 (2019).

41. Stortz, J. F. et al. Formin-2 drives polymerisation of actin filaments enabling segregation of apicoplasts and cytokinesis in Plasmodium falciparum. eLife 8, e49030 (2019).

42. Angrisano, F. et al. A GFP-actin reporter line to explore microfilament dynamics across the malaria parasite lifecycle. Mol. Biochem. Parasitol. 182, 93–6 (2012).

43. Bane, K. S. et al. The Actin Filament-Binding Protein Coronin Regulates Motility in Plasmodium Sporozoites. PLoS Pathog 12, e1005710 (2016).

44. Yahata, K., et al. Gliding motility of Plasmodium merozoites. Proc. Natl. Acad. Sci. U.S. A 118, e2114442118 (2021).

45. Ripp, J. et al. Malaria parasites differentially sense environmental elasticity during transmission. EMBO Mol Med 13, e13933 (2021).

46. Lettermann, L., Frischknecht, F. & XXX. TBA. Nat Phys (2025).

47. Amino, R. et al. Host cell traversal is important for progression of the malaria parasite through the dermis to the liver. Cell Host Microbe 3, 88–96 (2008).

48. Amino, R. et al. Quantitative imaging of Plasmodium transmission from mosquito to mammal. Nat. Med. 12, 220–4 (2006).

49. Munter, S. et al. Plasmodium sporozoite motility is modulated by the turnover of discrete adhesion sites. Cell Host Microbe 6, 551–62 (2009).

50. Briquet, S., Gissot, M. & Silvie, O. A toolbox for conditional control of gene expression in apicomplexan parasites. Mol Microbiol 117, 618–631 (2022).

51. Boyer, L. A. & Peterson, C. L. Actin-related proteins (Arps): conformational switches for chromatin-remodeling machines? BioEssays 22, 666–672 (2000).

52. Kast, D. J. & Dominguez, R. Arp you ready for actin in the nucleus? EMBO J 30, 2097–2098 (2011).

53. Andenmatten, N. et al. Conditional genome engineering in Toxoplasma gondii uncovers alternative invasion mechanisms. Nat Methods 10, 125–127 (2013).

54. Li, W. et al. A splitCas9 phenotypic screen in Toxoplasma gondii identifies proteins involved in host cell egress and invasion. Nat Microbiol 7, 882–895 (2022).

55. Biswas, A. et al. Paradoxical Effects of Substitution and Deletion Mutation of Arg56 on the Structure and Chaperone Function of Human αB-Crystallin. Biochemistry 46, 1117–1127 (2007).

56. Islam, M. M., Yohda, M., Kidokoro, S. & Kuroda, Y. Crystal structures of highly simplified BPTIs provide insights into hydration-driven increase of unfolding enthalpy. Sci Rep 7, 41205 (2017).

57. Andreadaki, M. et al. Genetic crosses and complementation reveal essential functions for the Plasmodium stage-specific actin2 in sporogonic development. Cell Microbiol 16, 751–67 (2014).

58. Schmitz, S. et al. Malaria parasite actin filaments are very short. J. Mol. Biol. 349, 113–25 (2005).

59. Lu, H., Fagnant, P. M. & Trybus, K. M. Unusual dynamics of the divergent malaria parasite PfAct1 actin filament. Proc. Natl. Acad. Sci. U.S.A. 116, 20418–20427 (2019).

60. Pražák, V., Vasishtan, D., Grünewald, K., Douglas, R. G. & Ferreira, J. L. Molecular architecture of glideosome and nuclear F-actin in Plasmodium falciparum. EMBO Rep 26, 1984–1996 (2025).

61. Sen, B. et al. Intranuclear Actin Regulates Osteogenesis. Stem Cells 33, 3065–3076 (2015).

62. Sen, B. et al. Intranuclear Actin Structure Modulates Mesenchymal Stem Cell Differentiation. Stem Cells 35, 1624–1635 (2017).

63. Kelpsch, D. J. & Tootle, T. L. Nuclear actin: from discovery to function. Anat Rec 301, 1999–2013 (2018).

64. Baum, J., Gilberger, T. W., Frischknecht, F. & Meissner, M. Host-cell invasion by malaria parasites: insights from Plasmodium and Toxoplasma. Trends Parasitol 24, 557–63 (2008).

65. Martinez, M. et al. Origin and arrangement of actin filaments for gliding motility in apicomplexan parasites revealed by cryo-electron tomography. Nat Commun 14, 4800 (2023).

66. Kan, A. et al. Quantitative analysis of Plasmodium ookinete motion in three dimensions suggests a critical role for cell shape in the biomechanics of malaria parasite gliding motility. Cell Microbiol 16, 734–50 (2014).

67. Trisnadi, N. & Barillas-Mury, C. Live In Vivo Imaging of Plasmodium Invasion of the Mosquito Midgut. mSphere 5, 10.1128/msphere.00692-20 (2020).

68. Beyer, K. et al. Limited Plasmodium sporozoite gliding motility in the absence of TRAP family adhesins. Malaria Journal 20, 430 (2021).

69. Braks, J. A., Franke-Fayard, B., Kroeze, H., Janse, C. J. & Waters, A. P. Development and application of a positive-negative selectable marker system for use in reverse genetics in Plasmodium. Nucleic Acids Res. 34, e39 (2006).

70. Janse, C. J. et al. High efficiency transfection of Plasmodium berghei facilitates novel selection procedures. Mol. Biochem. Parasitol. 145, 60–70 (2006).

71. Klug, D., Mair, G. R., Frischknecht, F. & Douglas, R. G. A small mitochondrial protein present in myzozoans is essential for malaria transmission. Open Biol 6, 160034 (2016).

72. Deligianni, E. et al. Critical role for a stage-specific actin in male exflagellation of the malaria parasite. Cell Microbiol 13, 1714–30 (2011).

73. Singer, M. et al. Zinc finger nuclease-based double-strand breaks attenuate malaria parasites and reveal rare microhomology-mediated end joining. Genome Biol 16, 249 (2015).

74. Schneider, C. A., Rasband, W. S. & Eliceiri, K. W. NIH Image to ImageJ: 25 years of Image Analysis. Nat Methods 9, 671–675 (2012).

75. Jumper, J. et al. Highly accurate protein structure prediction with AlphaFold. Nature 596, 583–589 (2021).

76. Sievers, F. et al. Fast, scalable generation of high-quality protein multiple sequence alignments using Clustal Omega. Mol. Syst. Biol. 7, 539 (2011).

