## Supplementary figures and tables for "Actin-related protein Alp1 governs malaria parasite motility and transmission"

**Table S1**: Primers used in the study (see separate Excel file)

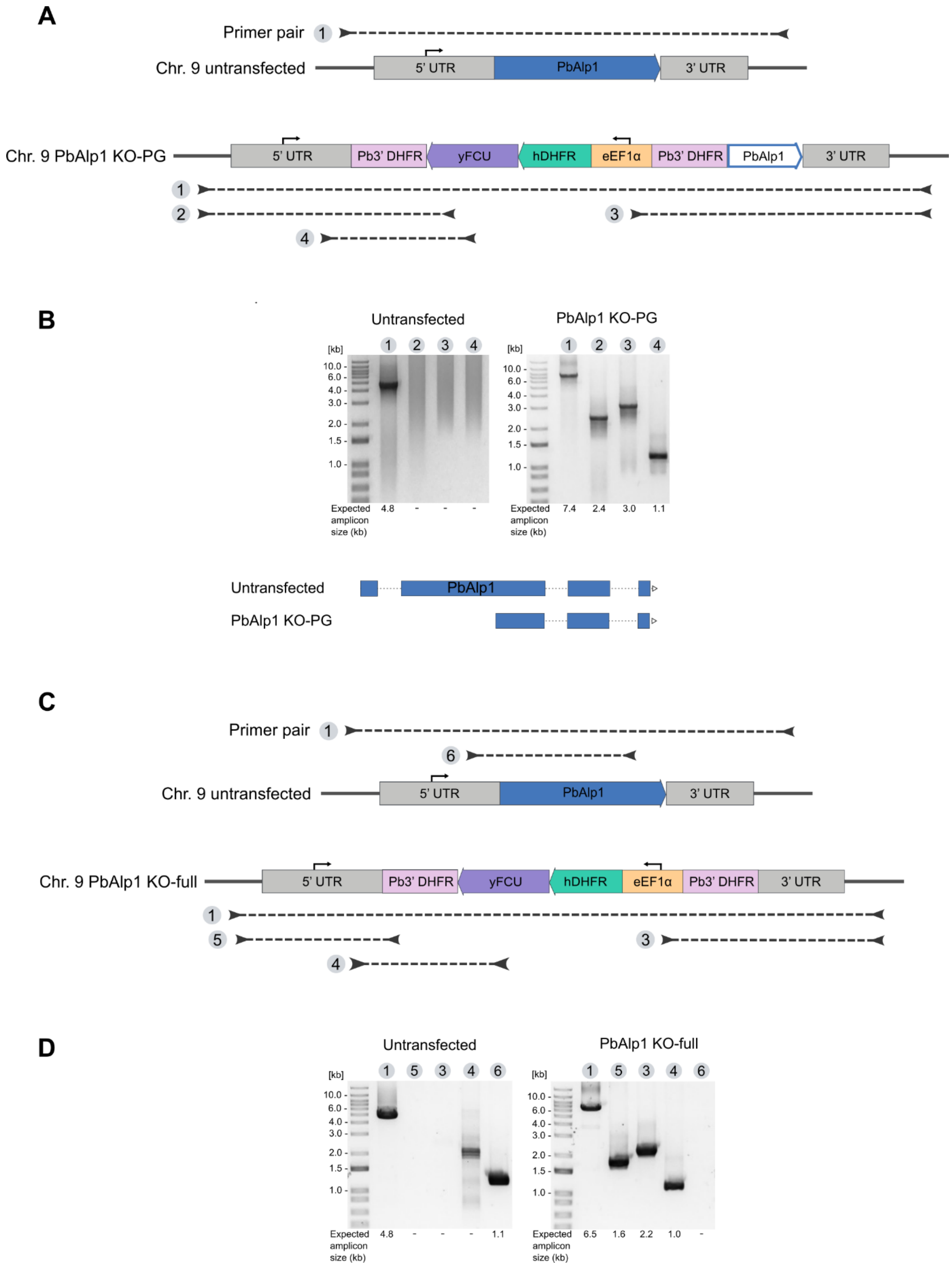


**Figure S1**: Generation and genotyping of Alp1 KO lines.
(A) Integration schemes and the genomic locus arrangement in both pre-transfected and transfected lines of PbAlp1 KO-PG. The orientation of the small arrows is indicative of the promoter, while the numbers and the dotted lines indicate the primer pairs used (see **Table S1**) and their coverage for genotyping. (B) Representative agarose gels of the selected genotyped parasite lines. Schemes below illustrate the structures of endogenous and the partially deleted *alp1*. (C) Integration schemes and the genomic locus arrangement in both

pre-transfected and transfected lines of PbAlp1 KO-full. The orientation of the small arrows is indicative of the promoter, while the numbers and the dotted lines indicate the primer pairs used (see **Table S1**) and their coverage for genotyping. (D) Representative agarose gels of the selected genotyped parasite lines.

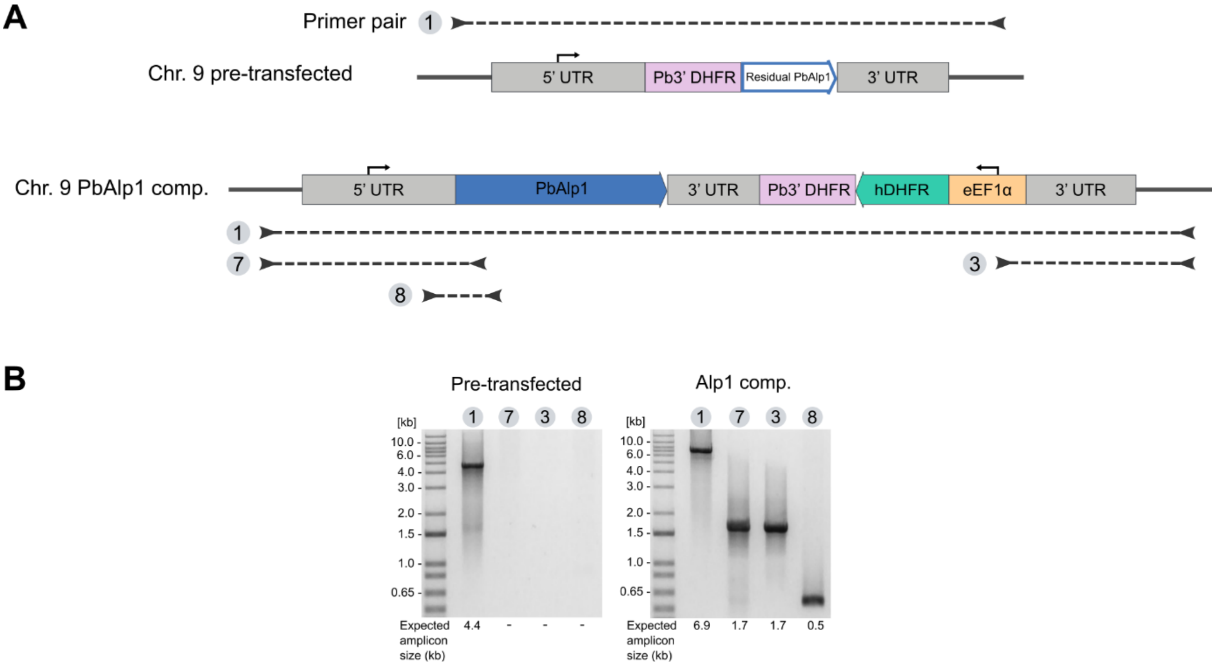


**Figure S2**: Generation and genotyping of PbAlp1 complementation line.
(A) Integration schemes and the genomic locus arrangement in both pre-transfected (PbAlp1 KO-PG) and transfected lines of PbAlp1 complementation. The orientation of the small arrows is indicative of the promoter, while the numbers and the dotted lines indicate the primer pairs used (see **Table S1**) and their coverage for genotyping. (B) Representative agarose gels of the selected genotyped parasite lines.

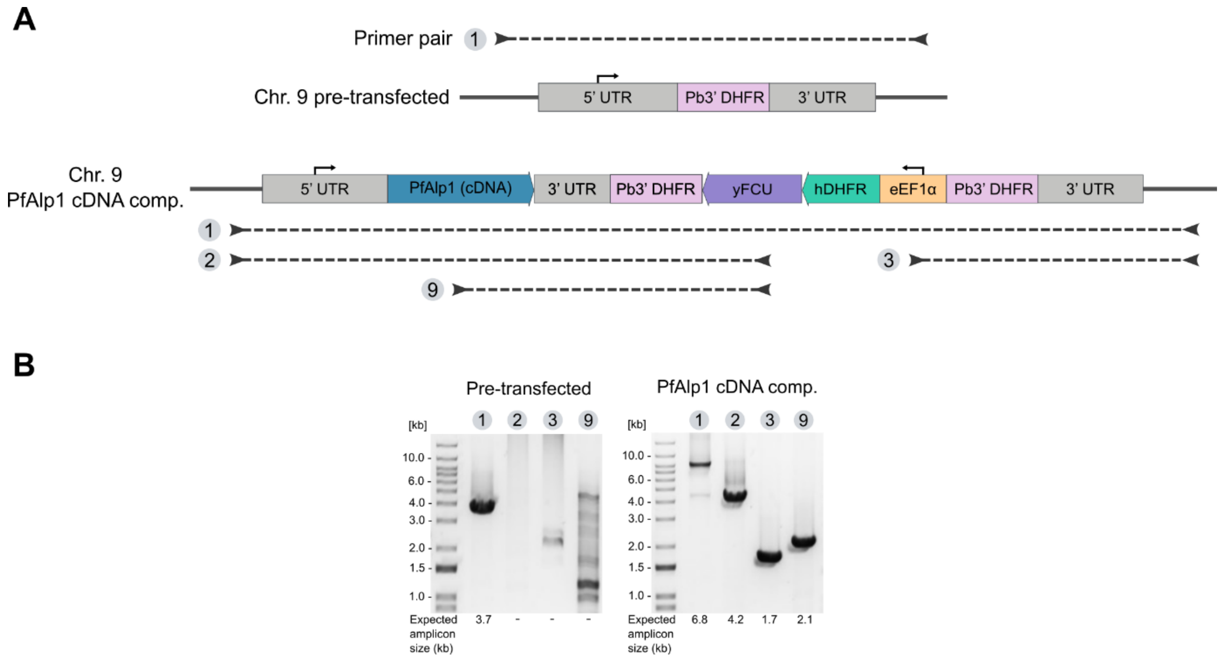


**Figure S3**: Generation and genotyping of PfAlp1 cDNA complementation line.
(A) Integration schemes and the genomic locus arrangement in both pre-transfected (PbAlp1 KO-full) and transfected lines of PfAlp1 cDNA complementation. The orientation of the small arrows is indicative of the promoter, while the numbers and the dotted lines indicate the primer pairs used (see **Table S1**) and their coverage for genotyping. (B) Representative agarose gels of the selected genotyped parasite lines.

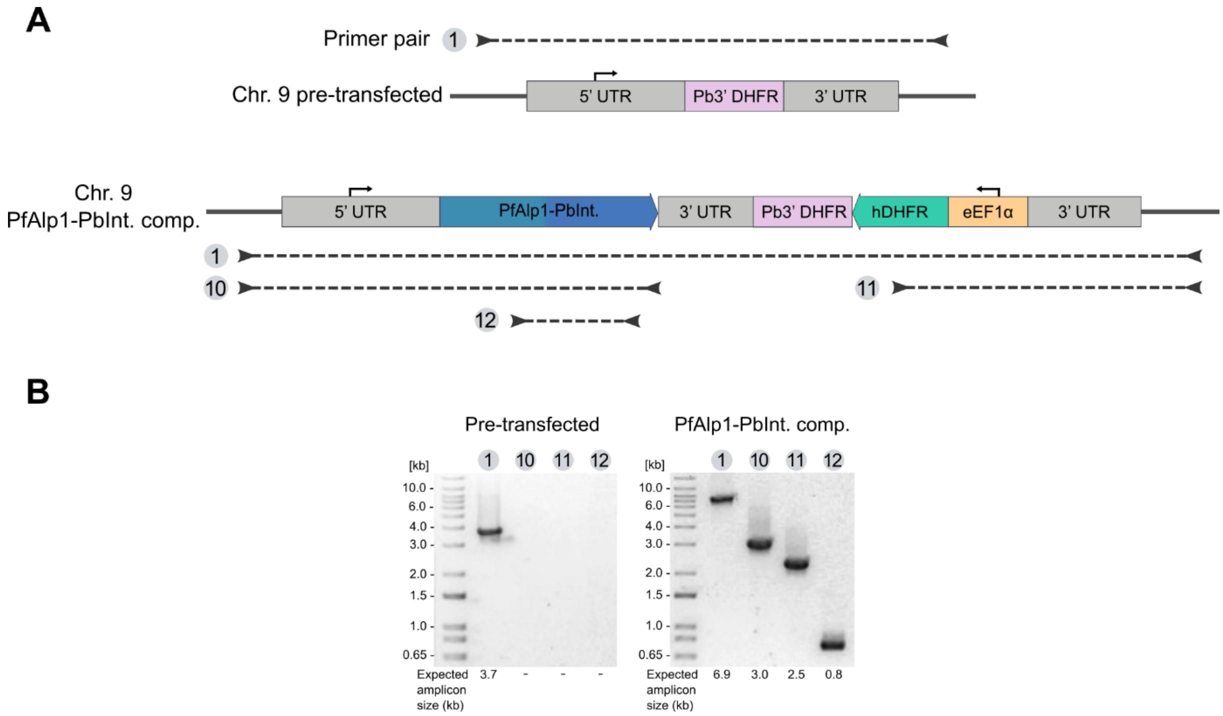


**Figure S4**: Generation and genotyping of PfEx.+PbIn. Alp1 complementation line. (A) Integration schemes and the genomic locus arrangement in both pre-transfected (PbAlp1 KO-full) and transfected lines of PfEx.+PbIn. Alp1 complementation. The orientation of the small arrows is indicative of the promoter, while the numbers and the dotted lines indicate the primer pairs used (see **Table S1**) and their coverage for genotyping. (B) Representative agarose gels of the selected genotyped parasite lines.

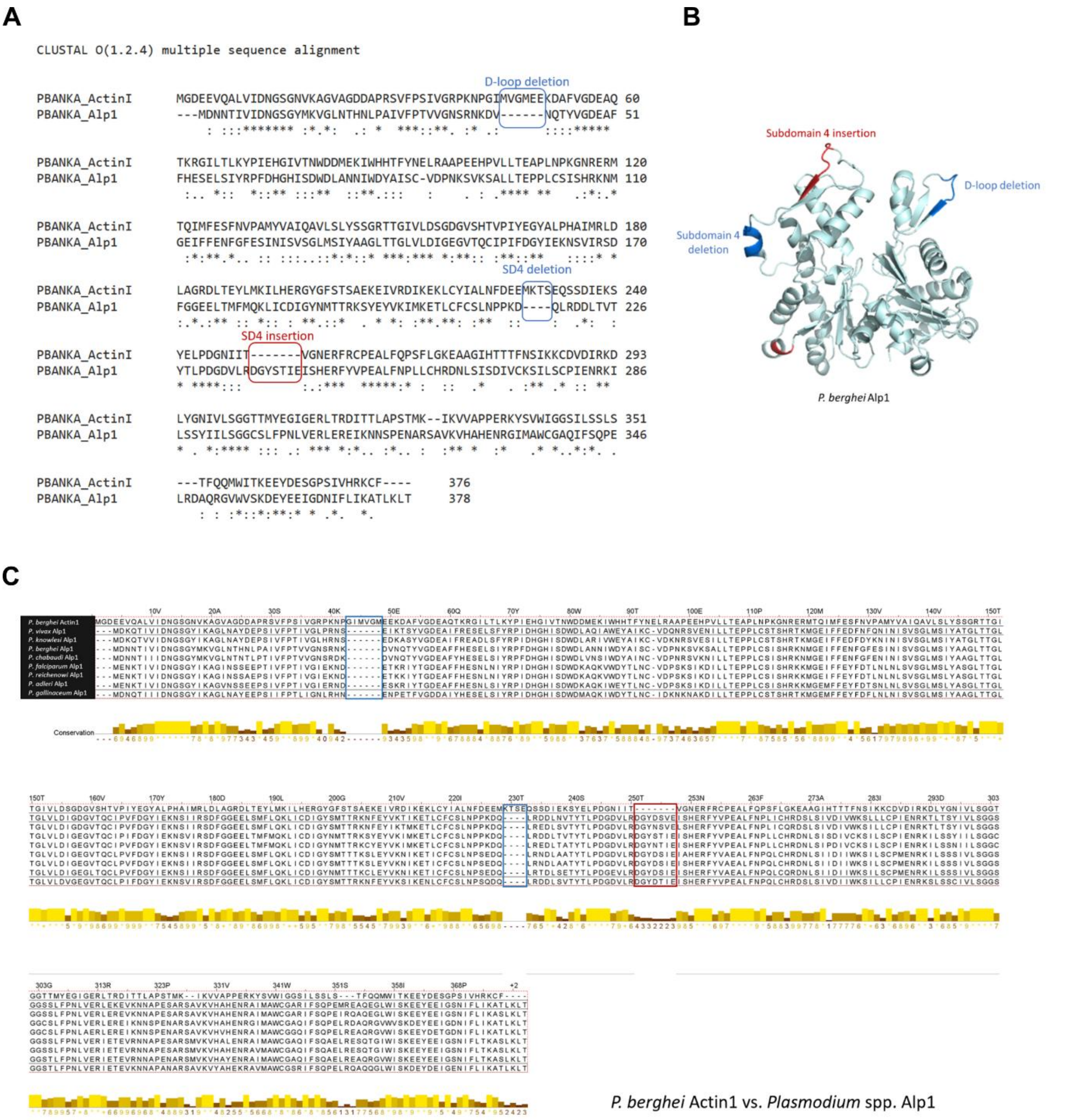


**Figure S5**: Sequence alignments of PbAlp1 with actin and Alp1 from other *Plasmodium* species.
(A) Protein sequence alignment of *P. berghei* actin 1 and Alp1, with the mutated insertion and deletion (InDel) regions indicated. For the actin insertion mutants (indicated in blue), amino acid residues from actin 1 were inserted into the Alp1 sequence: MVGMEE in the D-loop and MKTS in subdomain 4. Two mutations were introduced in the second deletion site in subdomain 4 (indicated in red): In one mutant, the amino acid residues in the Alp1 sequence (DGYSTIE) were deleted, while in the other, these residues were substituted with alanine. D-loop = DNase-I binding loop; SD = subdomain. (B) Predicted structure of PbAlp1 with InDel sites compared to Pbactin 1. Structure prediction was performed using AlphaFold2 (Jumper et al. 2021). (C) Protein sequence of Pbactin 1 aligned with Alp1 sequences from different *Plasmodium* species. Several unique insertions (red frame) and deletions (blue frames) are present in Alp1 in different *Plasmodium* species, compared to actin 1 in *P. berghei*. These InDels are highly conserved across species. The top row shows Pbactin 1, followed by Alp1 sequences from other *Plasmodium* species. All sequence alignments were conducted using Clustal Omega [76].

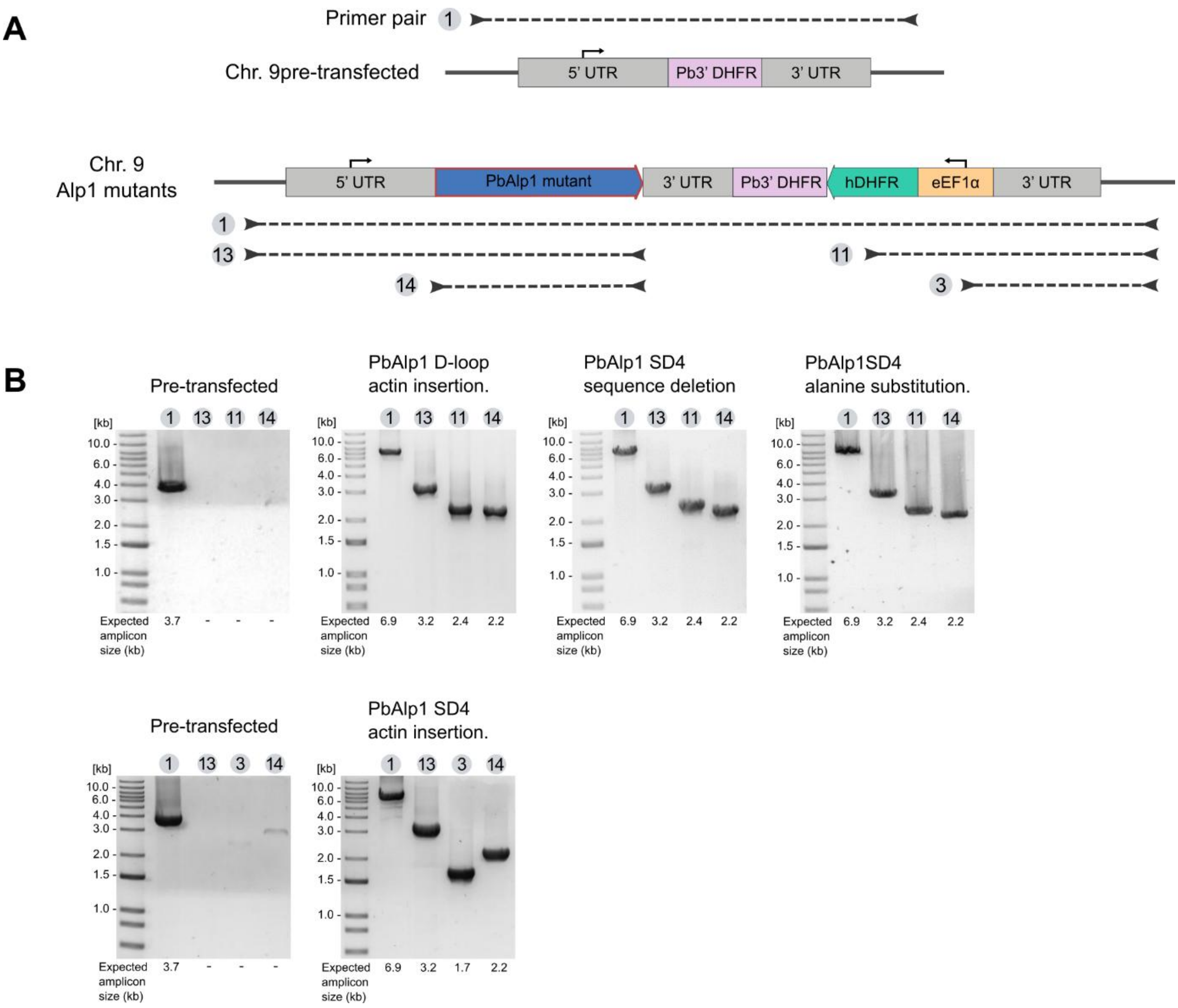


**Figure S6**: Generation and genotyping of Alp1 InDel mutant lines.
(A) Integration schemes and the genomic locus arrangement in both pre-transfected (PbAlp1 KO-full) and transfected lines of Alp1 InDel mutants. The orientation of the small arrows is indicative of the promoter, while the numbers and the dotted lines indicate the primer pairs used (see **Table S1**) and their coverage for genotyping. (B) Representative agarose gels of the selected genotyped parasite lines from each InDel mutant.

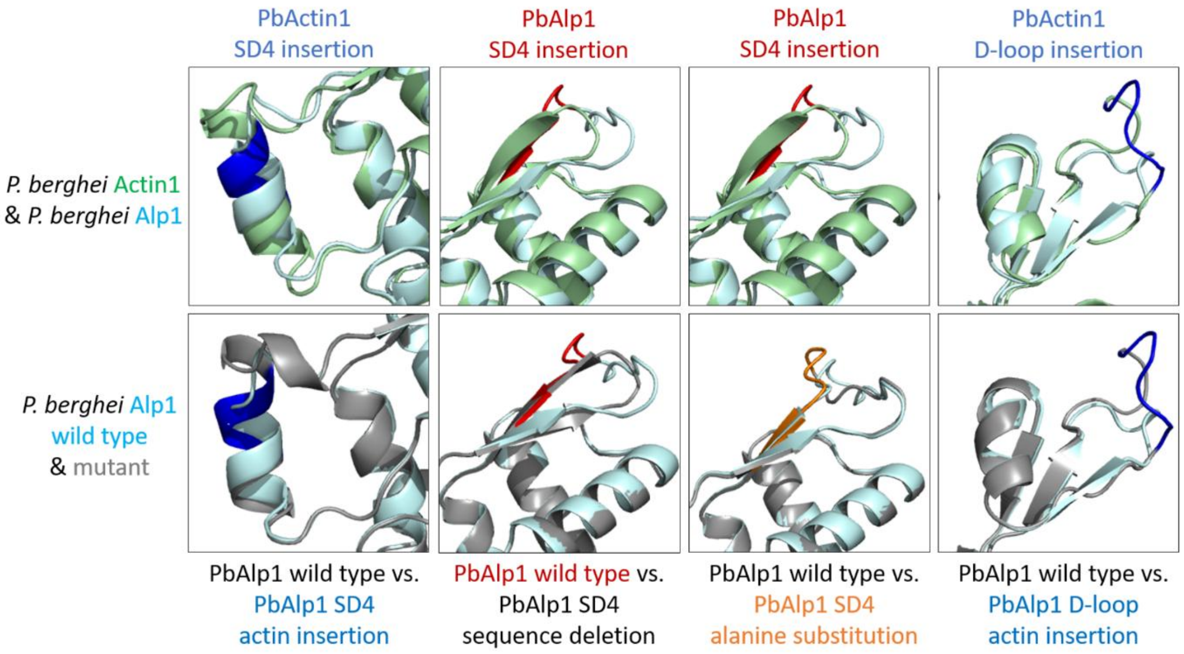


**Figure S7**: Predicted structural effects of mutations in Alp1.
In the top row, the enlarged images of the predicted Alp1 mutation sites taken from alignment of Pbactin 1 (green) and PbAlp1 (light blue) (**Figure S8**) are shown. Insertions are marked in dark blue or red as indicated by the description. The two middle panels with the PbAlp1 subdomain 4 (SD4) insertion are identical as two different mutations were generated at this site. In the bottom row, the mutated Alp1 sites (grey) are aligned with the predicted wild type (light blue) structures. The actin insertions are marked in blue and the alanine substitution in orange. All structure predictions were performed with AlphaFold2 and alignments were performed with open source PyMOL. As the top and bottom pictures were positioned and enlarged manually, minor differences in the size and tilt of the depicted protein regions could not be prevented. The mutations had no major effects on the predicted overall structures.

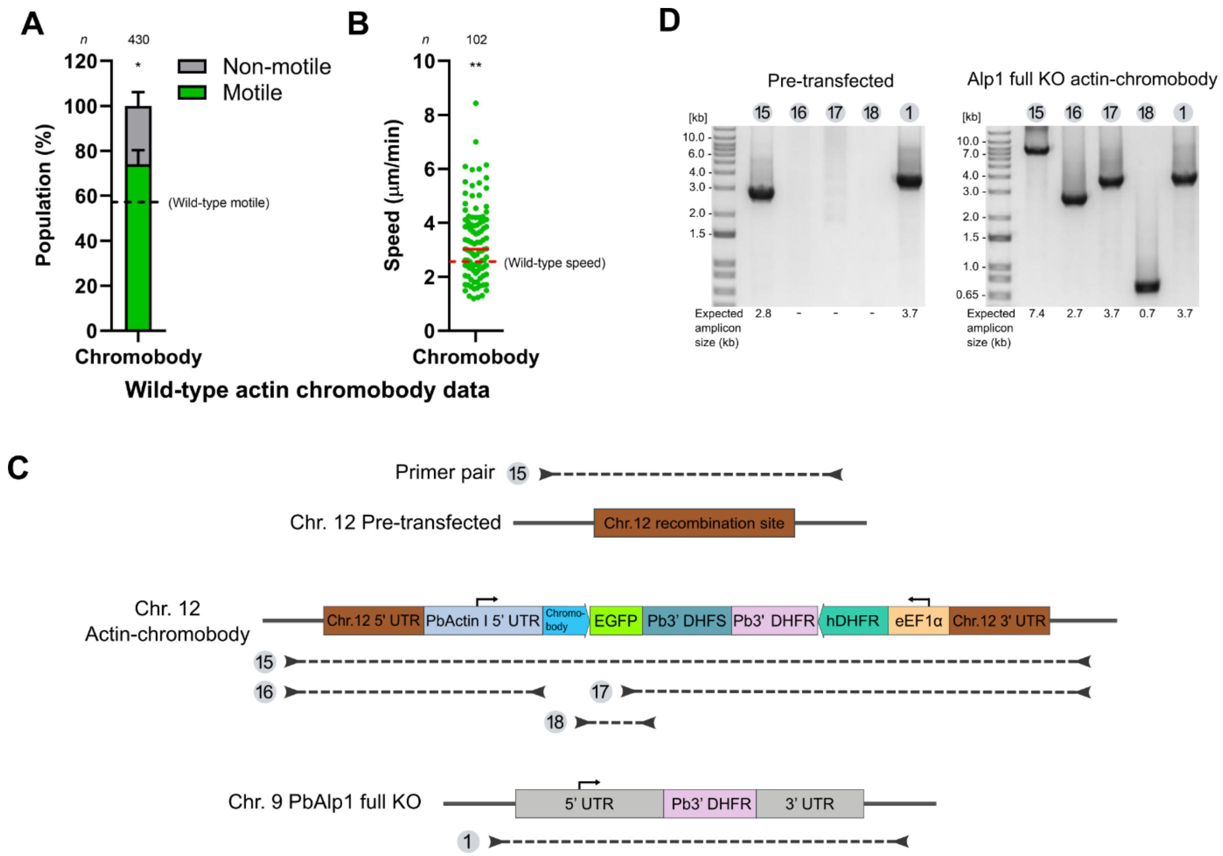


**Figure S8**: Impact of the actin chromobody on ookinete motility and generation and genotyping of the Alp1 KO actin chromobody line.
(A, B) Quantification of ookinete motility and speeds in the “wild-type” (i.e. no changes made in other loci) expressing actin chromobody line indicated that the chromobody has no negative effect on ookinete motility. The dotted line indicates the wild-type (no chromobody) data as in **Figure 2B-C**. Statistical analysis represents a level of significance against wild-type motile ookinetes and speeds (in A, Fisher's exact test, * $p<0.05$; in B, Mann-Whitney test for the speed, ** $p<0.01$). (C) Integration schemes and the genomic locus arrangement in both pre-transfected and transfected lines of Alp1KO-actin chromobody. The orientation of the small arrows is indicative of the promoter, while the numbers and the dotted lines indicate the primer pairs used (see **Table S1**) and their coverage for genotyping. (D) Representative agarose gels of the selected genotyped parasite lines. To ensure the absence of Alp1 in the transfected line, the corresponding locus was additionally genotyped.

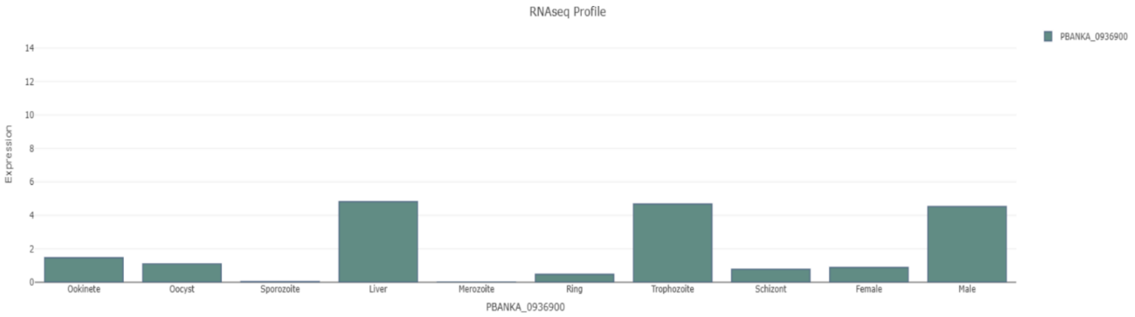


**Figure S9**: RNASeq profile of *P. berghei alp1* (PBANKA_0936900) throughout the life cycle. The bar graph, taken from the SPOT analysis (https://frischknechtlab.shinyapps.io/SPOT/) (Farr et al., 2021; Howick et al., 2019), shows that the *alp1* gene is highly expressed at the liver, trophozoite and male gamete stages. Meanwhile, it is less expressed in highly motile ookinetes and sporozoites.
