## Supplementary material for "Actin-related protein Alp1 governs malaria parasite motility and transmission": Table S1

**List of cloning primers**

| Line | Vector | Sequence | Purpose |
|---|---|---|---|
| Alp1 full KO | Pb262 | GATCGCGGCCGCTATGTGCAACTTTACCTAAAGG | PbAlp1 5' UTR amplification |
| | | GATCGATATCTCGTATAAGGTTATGTATGTTTAGTTATTTAATATTATG | |
| | | GCTAGGGCCCTTCAATTTATCTTTCCCGTTTTTTTTTC | PbAlp1 3' UTR amplification |
| | | ATGCGTTTAAACCGTACACACAAATATATGTGTATAC | |
| Alp1 comp | Pb238 | ATTCCGCGGACAATACGGTTTATTGGGG | PbAlp1 5' UTR + ORF amplification |
| | | ATAGGATCCCTAAGTTAGCTTTAGAGTTGCC | |
| | | AAACCTAGGTTCAATTTATCTTTCCCGTTTTTTTTTC | PbAlp1 3' UTR-outer amplification |
| | | ATAGGTACCCTTTCTTTTTAAATATAGGTAAAAGGTC | |
| | | AAAGGATCCGTTCAATTTATCTTTCCCGTTTTTTTTTC | PbAlp1 3' UTR-inner amplification |
| | | TTTGATATCATACTATGAATAAAATAAAAATGGAAAATC | |
| Alp1 Pf comp | Pb262 | GATCGAATTCTTCAATTTATCTTTCCCGTTTTTTTTTC | PbAlp1 3' UTR-inner amplification |
| | | CGATGATATCGTGAGAATAATCAAACTAGTGC | |
| | | CAGTGCGGCCGCATGGAGAACAAGACAATAGTAATTGATAAC | PfAlp1 ORF amplification |
| | | CAGTGAATTCTTAAGTAAGTTTTAGAGATGCCTTTG | |
| | | GCATGCGCGCTATGTGCAACTTTACCTAAAGG | PbAlp1 5' UTR amplification |
| | | ATATGCGGCCGCTCGTATAAGGTTATGTATGTTTAGTTATTTAATATTATG | |
| | | ATGCGTTTAAACCGTACACACAAATATATGTGTATAC | PbAlp1 3' UTR-outer amplification |
| | | GCATGGCCGGCCTTCAATTTATCTTTCCCGTTTTTTTTTC | |
| Alp1 Pf-PbInt comp | Pb238 | GCATGCGCGCTATGTGCAACTTTACCTAAAGG | PfAlp1 ORF exon 1 amplification |
| | | CAATAATTTTTTTACCTGAACCGTTATCAATTACTATTGTCTTG | |
| | | GATAACGGTTCAGGTAAAAAAATTATTGCATGTGCAGG | PbAlp1 ORF intron 1 amplification |
| | | CTTTATATATCCTATATATTGAGAAATTAATAAAAATAGCGAAATTGAGG | |
| | | CTCAATATATAGGATATATAAAGGCGGGAATAAATTCAAGTG | PfAlp1 ORF exon 2 amplification |
| | | TTAAAATTACCTTGACCATCGAACGGG | |
| | | GTTCGATGGTCAAGGTAATTTTAAATGAACAAACTGTGTCTATC | |

| | | | |
|---|---|---|---|
| | | GAGCATGAACCTGAAGATTAAAAAGTAGAAGCATCAAAG | PbAlp1 ORF intron 2 amplification |
| | | CTTTTTAATCTTCAGGTTCATGCTCTTGAGAACAGAG | PfAlp1 ORF exon 3 amplification |
| | | GTTTTATTTTTTTTCTTACCTTTGTTAAAAAAATGTTGGATCCTATTTC | |
| | | CATTTTTTTAACAAAGGTAAGAAAAAAAATAAAACAAATAGAGTATACAAAACG | PbAlp1 ORF intron 3 + PfAlp1 ORF exon 4 amplification |
| | | CAGATCTTTAAGTAAGTTTTAGAGATGCCTATAAAAAAAATATTAAAATAACCAAAGTCAATAAAATTG | |
| Alp1 mutant E1 D-loop ins | Pb238 | ACTGCGCGCGGACAATACGGTTTATTG | PbAlp1 5' ORF + Act1 D-loop amplification |
| | | TTCTTCCATCCCTACCATTACATCTTTGTTTCTCGAATTTCC | |
| | | GTAATGGTAGGGATGGAAGAAAATCAAACATATGTGGGCG | PbAlp1 3' ORF + Act1 D-loop amplification |
| | | GACTGGATCCCTAAGTTAGCTTTAGAG | |
| Alp1 mutant E2 SD4 del | Pb238 | ACTGCGCGCGGACAATACGGTTTATTG | PbAlp1 5' ORF + SD4 deletion amplification |
| | | ATGTGAAATTCTTAAAACATCCCCATCAGGTAATG | |
| | | TGTTTTAAGAATTTCACATGAACGATTTTATGTTC | PbAlp1 3' ORF + SD4 deletion amplification |
| | | GACTGGATCCCTAAGTTAGCTTTAGAG | |
| Alp1 mutant E3 SD4 Ala | Pb238 | ACTGCGCGCGGACAATACGGTTTATTG | PbAlp1 5' ORF + SD4 alanine insertion amplification |
| | | ATGTGAAATTGCAGCTGCTGCAGCTGCTGCTCTTAAAACATCCCCATCAGG | |
| | | GCAGCAGCTGCAGCAGCTGCAATTTCACATGAACGATTTTATGTTCC | PbAlp1 3' ORF + SD4 alanine insertion amplification |
| | | GACTGGATCCCTAAGTTAGCTTTAGAG | |
| Alp1 mutant E4 SD4 Act | Pb238 | ACTGCGCGCGGACAATACGGTTTATTG | PbAlp1 5' ORF + SD4 Act1 insertion amplification |
| | | AATCATCTCTTAATTGAGATGTTTTCATATCTTTTGGAGGGTTTAACG | |
| | | ATGAAAACATCTCAATTAAGAGATGATTTAACAGTTACATATAC | PbAlp1 3' ORF + SD4 Act1 insertion amplification |
| | | GACTGGATCCCTAAGTTAGCTTTAGAG | |
| Alp1 full KO - chromobody | Yee et al. (2022) | | |

### List of genotyping primer pairs

| Primer pair | Direction | Primer sequence (5' - 3') |
|---|---|---|
| 1 | Forward | GTTCTAGTGTAACTTGAATTGGAAC |
| | Reverse | GCTACAATGCATAAAGGATACG |
| 2 | Forward | GTTCTAGTGTAACTTGAATTGGAAC |
| | Reverse | GAGAGGTGTTAAGCCAGAG |
| 3 | Forward | GGGGTGAGCATTTAAAGC |
| | Reverse | GCTACAATGCATAAAGGATACG |
| 4 | Forward | TATTGGACATTTGGGGGG |
| | Reverse | TAATTCAAAGGGACGAGG |
| 5 | Forward | TTCTTTGATATGCTTTTTGTTCTTTC |
| | Reverse | TCCTTCAATTTCGATGGGTAC |
| 6 | Forward | TATTGGACATTTGGGGGG |
| | Reverse | ATCTCTATGACATAAAAGTGG |
| 7 | Forward | TTCTAGTGTAACTTGAATTGGAAC |
| | Reverse | ACTTATTTGCCTGCACATGC |
| 8 | Forward | TATTGGACATTTGGGGGG |
| | Reverse | CTTCGTCGCCCACATATG |
| 9 | Forward | GTTCAGATTTCGGTGGAGAAG |
| | Reverse | GAGAGGTGTTAAGCCAGAG |
| 10 | Forward | GTTCTAGTGTAACTTGAATTGGAAC |
| | Reverse | GTTTTATTTTTTTTCTTACCTTTGTTAAAAAAATGTTGGATCCTATTTC |
| 11 | Forward | GTGGAGGTTCCTTGAGTTC |
| | Reverse | GCTACAATGCATAAAGGATACG |
| 12 | Forward | GTTCAGATTTCGGTGGAGAAG |
| | Reverse | GAGAGGTGTTAAGCCAGAG |
| 13 | Forward | GTTCTAGTGTAACTTGAATTGGAAC |

| | | |
|---|---|---|
| | Reverse | GACTGGATCCCTAAGTTAGCTTTAGAG |
| 14 | Forward | ACTGCGCGCGGACAATACGGTTTATTG |
| | Reverse | GACTGGATCCCTAAGTTAGCTTTAGAG |
| 15 | Forward | GAAGACAACCAAGACGATCTTG |
| | Reverse | GTAGCTCGAGGATGATTTAGAATCTTTATATGCACCTATGC |
| 16 | Forward | GAAGACAACCAAGACGATCTTG |
| | Reverse | GCTCTAGATTTAATTTTTTTTTTAAGTATATGAGTATATATATGTGTGTAAAAATTTATATTAAATATGC |
| 17 | Forward | CACAACATCGAGGACGGC |
| | Reverse | GTAGCTCGAGGATGATTTAGAATCTTTATATGCACCTATGC |
| 18 | Forward | ATATGCGGCCGCATGGTGAGCAAGGGCGAG |
| | Reverse | AGCCCTAGGTTACTTGTACAGCTCGTCC |
